# Structural insights into *Pseudomonas aeruginosa* lysine-specific uptake mechanism for extremely low pH regulation

**DOI:** 10.1101/2024.05.17.594718

**Authors:** Deniz Bicer, Rei Matsuoka, Aurélien F. A. Moumbock, Preethi Sukumar, Harish Cheruvara, Andrew Quigley, Els Pardon, Jan Steyaert, Peter J.F. Henderson, Martin Caffrey, Julia J. Griese, Emmanuel Nji

**Affiliations:** Department of Cell and Molecular Biology, Uppsala University, 751 24 Uppsala, Sweden; OMass Therapeutics Ltd, Building 4000, Chancellor Court, John Smith Drive, ARC Oxford, OX4 2GX, UK; Institute of Pharmaceutical Sciences, Albert-Ludwigs-Universität Freiburg, Freiburg, Germany; Astbury Centre for Structural Molecular Biology, University of Leeds, Leeds LS2 9JT, UK; Membrane Protein Laboratory, Diamond Light Source Ltd., Research Complex at Harwell, Didcot OX11 0DE, UK; VIB-VUB Center for Structural Biology, VIB, Pleinlaan 2, 1050 Brussels, Belgium; Structural Biology Brussels, Vrije Universiteit Brussel, Pleinlaan 2, 1050 Brussels, Belgium; Schools of Medicine and Biochemistry & Immunology, Trinity College, Dublin D02 R590, Ireland; Department of Parasitology and Microbiology, Centre for Research in Infectious Diseases, P.O. Box 13591, Yaoundé, Cameroon; BioStruct-Africa, Alfred Nobels Allé 37C, 1001-14152 Huddinge, Sweden; BioStruct-Africa, 10 Link Road, Kwame Nkrumah University of Science and Technology (KNUST), Kumasi, Ghana

## Abstract

Under conditions of extremely low pH, in addition to transporting lysine, bacterial lysine-specific permease (LysP) interacts with the transcriptional regulator CadC to upregulate *cadBA* operon expression. *cadBA* encodes CadA, which decarboxylates lysine to cadaverine, and CadB, which exports cadaverine to the environment to reduce acidity. This process is crucial for survival of pathogenic bacteria in their hosts. Here, we report the inward-occluded (3.2 – 5.3 Å) cryo-EM structure of *Pseudomonas aeruginosa* LysP bound to L-lysine and in complex with a nanobody. L-lysine is coordinated by hydrophobic stacking, cation-π interactions and hydrogen bonding mostly with polar uncharged LysP residues. LysP reconstituted into liposomes showed robust and specific transport of L-lysine with the transport being inhibited by L-4-thialysine (S-2-aminoethyl-L-cysteine). These findings inform our understanding of the specific recognition, inhibition, and transport mechanism of L-lysine by LysP, which will have important ramifications for the design of antibiotics to target bacterial LysP.

## Main

Antibiotic resistance is a significant global public health threat that can have severe consequences, including increased morbidity, mortality, and healthcare costs^1–4^. The number of deaths due to antibiotic-resistant bacteria has been increasing in the European Union and the European Economic Area, with over 25,000 deaths reported in 2007^5^, 33,000 in 2015^5^ and 38,710 in 2019^6^. The increase in deaths in 2019 was associated with an estimated 865,767 infections of selected antibiotic-resistant bacteria, leading to 1,101,288 total disability-adjusted life-years (DALYs)^6^. In the United States, antibiotic resistance is also a significant public health threat, with more than 2.8 million people infected and over 35,000 deaths reported each year due to antibiotic-resistant infections^7^. Furthermore, according to the World Health Organization (WHO), if appropriate measures are not taken to address antibiotic resistance, the number of deaths due to antibiotic-resistant infections is predicted to rise significantly, with a projected 10 million deaths per year globally by 2050^8^. This threat further emphasizes the importance of developing strategies to address antibiotic resistance, such as promoting the appropriate use of antibiotics in medicine, veterinary and agriculture, promoting the use and development of vaccines and tools for early diagnosis of bacterial infections, educating the community about the danger of antibiotic resistance, and investing in research to find alternative treatments for bacterial infections. Here, we investigate the structure-function relationship of the lysine-specific permease (LysP), a protein involved in the transport of lysine, an essential amino acid, across the inner cell membrane in *P. aeruginosa*. Under conditions of extremely low pH, as in the mammalian stomach, LysP plays a crucial role in the survival of bacteria by interacting with the transcriptional regulator CadC to upregulate the expression of the *cadBA* operon^9–11^. The *cadBA* operon encodes two proteins, CadA and CadB, which, respectively, are involved in the decarboxylation of lysine to cadaverine and the export of the alkaline product cadaverine to the environment to neutralize the proton and thus reduce the acidity (Fig. 1a)^9–13^. The decarboxylation of lysine consumes a proton which contributes to the proton gradient that facilitates cadaverine transport^14^.

**Fig. 1.**
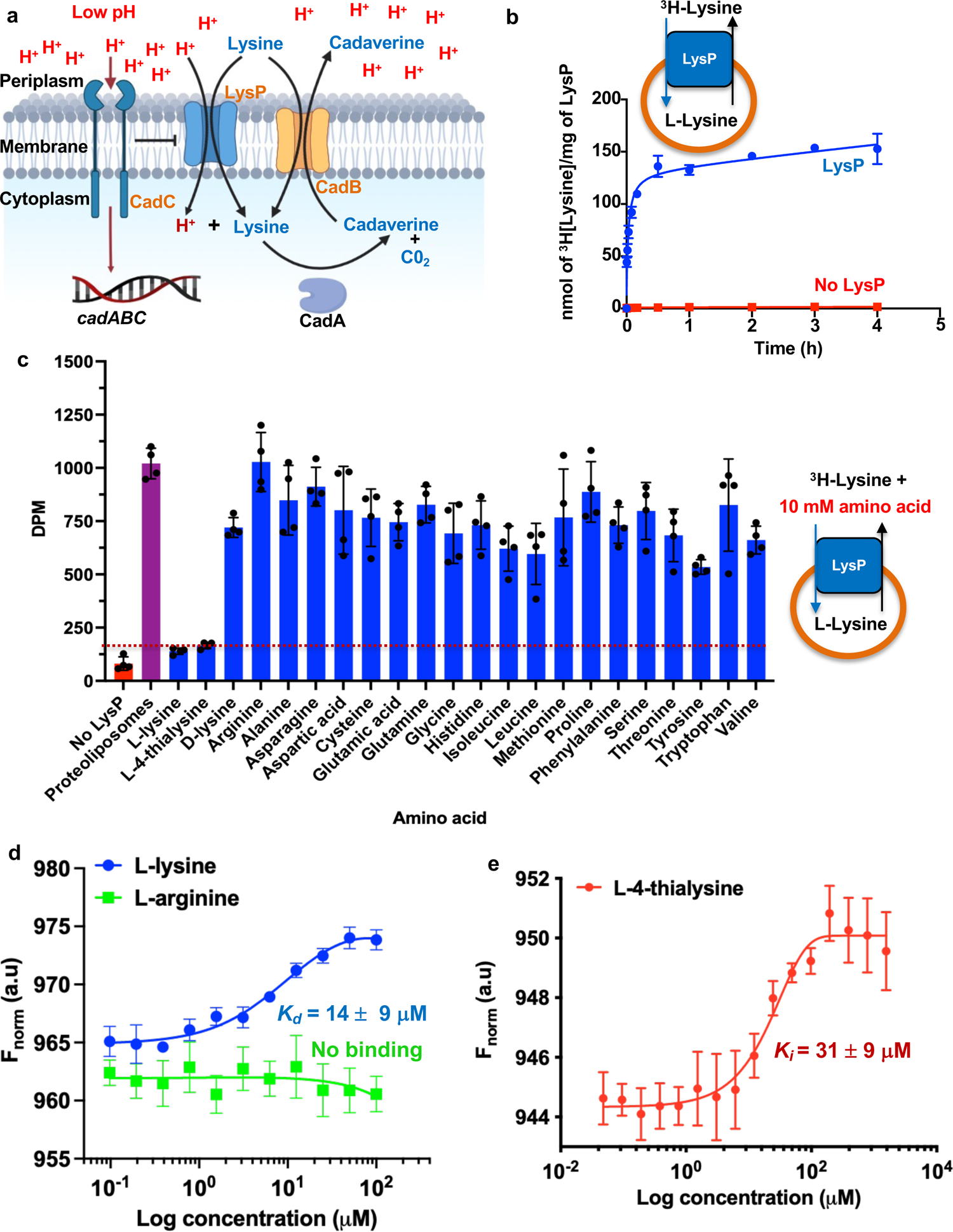
Functional characterization of LysP. **a,** Schematic representation of bacterial lysine-specific transport mechanism and its role in extremely low pH regulation. **b,** [^3^H] L-lysine counterflow uptake by reconstituted LysP. Blue shows transport into proteoliposomes containing LysP and red shows transport into liposomes with no LysP (control). The mean ± s.e.m. of the fit is the average from three independent experiments for liposomes and two for liposomes **c,** Inhibition of [^3^H]-L-lysine transport into proteoliposomes reconstituted with LysP. The inhibition was carried out in the presence of 10 mM of the different cold substrates. The mean ± s.e.m. of the fit is the average from four independent experiments **d,** Binding of L-lysine and L-arginine to LysP by microscale thermophoresis (MST). The mean ± s.e.m. of the fit is the average from three independent experiments **e,** Binding of thialysine to LysP by MST. The mean ± s.e.m. of the fit is the average from three independent experiments.

This process is important for bacterial survival in the host, as it allows the bacteria to maintain a neutral pH environment, which is necessary for growth^9–11^. The LysP protein from *Pseudomonas aeruginosa* is similar in sequence to LysP from other enteric pathogens (Extended Data Fig. 1a & b), which cause a significant disease burden worldwide, contributing to more than 96 million DALYs globally^15^.

The structure and mechanism of LysP from *P. aeruginosa* (hereafter referred to as LysP), a pathogenic bacterium, is of great interest as this organism is on the WHO list of priority pathogens, known as ESKAPE (*Enterococcus faecium*, *Staphylococcus aureus*, *Klebsiella pneumoniae*, *Acinetobacter baumannii*, *Pseudomonas aeruginosa*, and *Enterobacter species*), for research and development of new antibiotics^16^. *P. aeruginosa* can cause infections in the urinary tract, respiratory system, soft tissue, bones and joints, and many systemic infections such as bacteremia and dermatitis, particularly in hospitalized patients with cystic fibrosis, severe burns, cancer and AIDS patients whose immune systems have been compromised^17–19^. Indeed, 10 % of all hospital-acquired infections are caused by *P. aeruginosa* which is difficult to treat because of a naturally high antibiotic resistance and have a mortality rate as high as 40 %^17–20^.

*P. aeruginosa* LysP is an integral membrane lysine specific transporter that belongs to the <u>a</u>mino acid, <u>p</u>olyamine and organo<u>c</u>ation (APC) transporter superfamily^21,22^. APC family transporters are present in all kingdoms of life and are vital in cell physiology and highly relevant in diseases, which include the human solute carrier (SLC) families^11^. APC family transporters are involved in nutrient uptake, elimination of toxic waste, and exchange of information and signals^21,22^.

While the crystal and/or cryo-EM structures of ApcT (Na^+^-independent amino acid transporter)^23^, AdiC (arginine-agmatine antiporter)^24–26^, GadC (glutamate-GABA antiporter)^27^, AgcS (Na^+^-alanine symporter)^28^, BasC (alanine-serine-cysteine exchanger)^29^, CCC (cation-chloride cotransporter)^30–37^, and KimA (K^+^/H^+^-symporter)^38^, along with decades of biochemical and biophysical studies have provided valuable insights into the selectivity and mode of action of these transporters, there is still much to learn about the structure-function relationship of this diverse family of transporters. For example, many amino acid transporters are known to exhibit promiscuous substrate recognition, meaning that they can transport multiple substrates with varying affinities^23,39^. The structural basis for this promiscuity is not well understood, and further research is necessary to elucidate the mechanisms of substrate recognition and transport. Here, we investigated the first transporter that specifically transports lysine from *P. aeruginosa* using single-particle cryo-EM and functional approaches. The structure of LysP bound to L-lysine and in complex with a nanobody, as well as accompanying functional data reveal useful molecular insights into how LysP, unlike the broad-spectrum amino acid transporters, utilizes a proton gradient to transport with high specificity L-lysine, and how it is inhibited by S-(2-aminoethyl)-L-cysteine (L-4-thialysine), which will open up avenues for structure-based antibiotics design to target bacterial LysP.

## Results

### Cryo-EM structure determination

Several attempts to obtain well-diffracting crystals of LysP failed^40,41^. To obtain stable protein for structure determination, camelid nanobodies were generated and screened against LysP (Extended Data Fig. 4a & b). The best binder Nb5755 improved the stability of LysP with an increase in melting temperature of 20 °C (Fig. 2a) and the complex was used for cryo-EM studies. The binding of Nb5755 to LysP was also evident by a peak shift during size exclusion chromatography (SEC) towards higher molecular weight (Fig. 2b and Extended Data Fig. 4b), as well as analysis on SDS PAGE of the SEC peak fraction of the complex which showed two bands corresponding to LysP and Nb5755 (Fig. 2c and Extended Data Fig. 4b). The LysP protein without nanobody in solution revealed predominantly an α-helical profile as judged by circular dichroism (Fig. 2d), and importantly, LysP was functional in transport assays (Fig. 1b), making the sample suitable for single-particle cryo-EM analysis. Particle averaging yielded a cryo-EM map with an overall resolution of ∼3.68 Å for the LysP-L-lysine-Nb5755 complex (Fig. 2e, 2f, Table 1 and Extended Data Fig. 2).

**Fig. 2.**
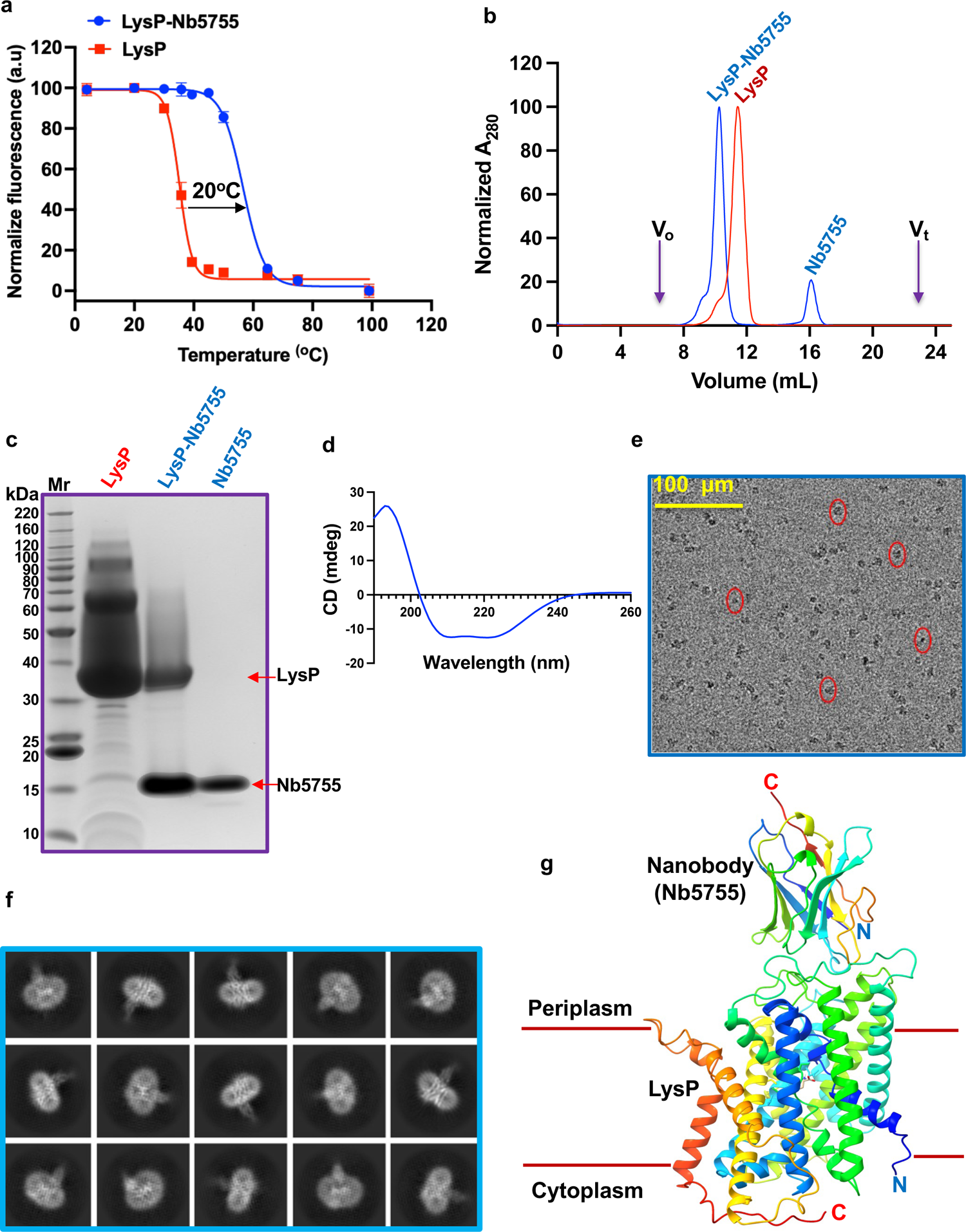
Biochemical and structural characterization of LysP. **a,** Green Fluorescent Protein Thermal Shift (GFP-TS) melting curve of LysP-GFP with Nb5755 (blue) and without Nb5755 (red). Each apparent *T_m_* (mean ± s.e.m. of the fit) is the average from three independent experiments **b,** Gel filtration chromatogram of LysP (red) and LysP-Nb5755 complex (blue). The small peak around 16 mL represents excess free Nb5755 (blue). n=1 for a single experiment **c,** SDS PAGE analysis of the peaks shown in **b**, lane 1 (free LysP), lane 2 (LysP-Nb5755 complex) and lane 3 (free Nb5755) and Mr (molecular weight standards). **d,** Circular dichroism spectrum of free LysP. n=3 for three independent experiments **e,** Example micrograph depicting extraction boxes (areas of interest extracted from the micrograph for analysis in red circles). **f,** Representative 2D classes of LysP-Nb5755. **g,** 3D Cryo-EM structure of LysP-Nb5755-L-lysine complex.

**Table 1.** Cryo-EM data collection, refinement and validation statistics.

|  | LysP-Nanobody 5755 complex<br>(EMDB-50053)<br>(PDB 9EYD) |
| --- | --- |
| **Data collection and processing** |  |
| Magnification | 130000 |
| Voltage (kV) | 300 |
| Electron exposure (e<sup>-</sup>/Å<sup>2</sup>) | 38.7 |
| Defocus range (μm) | -0.8 to -2.4 |
| Pixel size (Å) | 0.6725 |
| Symmetry imposed | None |
| Initial particle images (no.) | 177424 |
| Final particle images (no.) | 78459 |
| Map resolution (Å) | 3.68 |
| FSC threshold | 0.143 |
| Map resolution range (Å) | 3.2 to 5.3 |
| **Refinement** |  |
| Initial model used (PDB code) | None |
| Model resolution (Å) | 3.7 |
| FSC threshold | 0.143 |
| Model resolution range (Å) | 3.2 to 4.1 |
| Map sharpening B factor (Å<sup>2</sup>) | -138 |
| Model composition |  |
| Non-hydrogen atoms | 4568 |
| Protein residues | 595 |
| Ligands | - |
| B factors (Å<sup>2</sup>) |  |
| Protein | 122.9 |
| Ligand | 85.8 |
| R.m.s. deviations |  |
| Bond lengths (Å) | 0.002 |
| Bond angles (°) | 0.475 |
| **Validation** |  |
| MolProbity score | 1.93 |
| Clashscore | 4.49 |
| Poor rotamers (%) | 3.02 |
| **Ramachandran plot** |  |
| Favored (%) | 95.06 |
| Allowed (%) | 4.94 |
| Disallowed (%) | 0 |

### Structure of LysP–L-Lysine-Nb5755 in an inward-occluded state

The structure of the LysP-Nb5755 complex was determined in the presence of L-lysine (Fig. 3, Fig. 4 and Extended Data Fig. 3a). LysP adopted an inward-occluded state in the transport cycle (Fig. 6b) with the nanobody binding to LysP on the periplasmic side (Fig. 2g). The LysP structure reveals 12 transmembrane helices with both C- and N-termini located in the cytoplasm as previously described^42^ (Fig. 2g, Fig. 3 and Extended Data Fig. 3b).

**Fig. 3.**
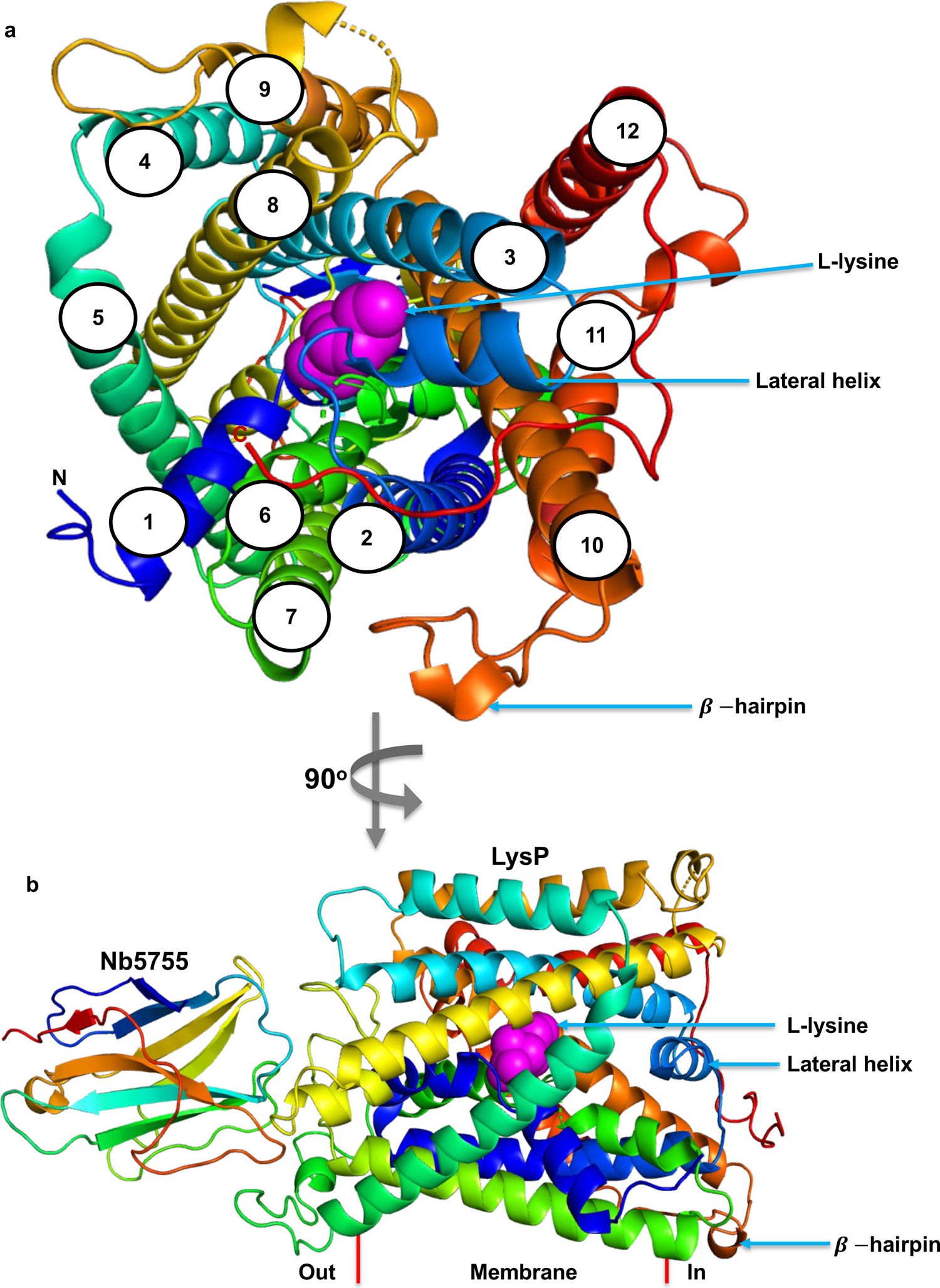
Structure of LysP-Nb5755-L-lysine complex. **a**, Cytoplasmic view of the LysP-Nb5755 structure in complex with L-lysine. LysP has 12 transmembrane helices with both C- and N-termini located in the cytoplasm. Helices 1 and 6 are broken in the middle, Helices 2 and 3 are linked by a lateral cytoplasmic helix. Helices 10 and 11 are linked by a β-hairpin. **b**, 90° rotation of the cytoplasmic view shown in **a**.

**Fig. 4.**
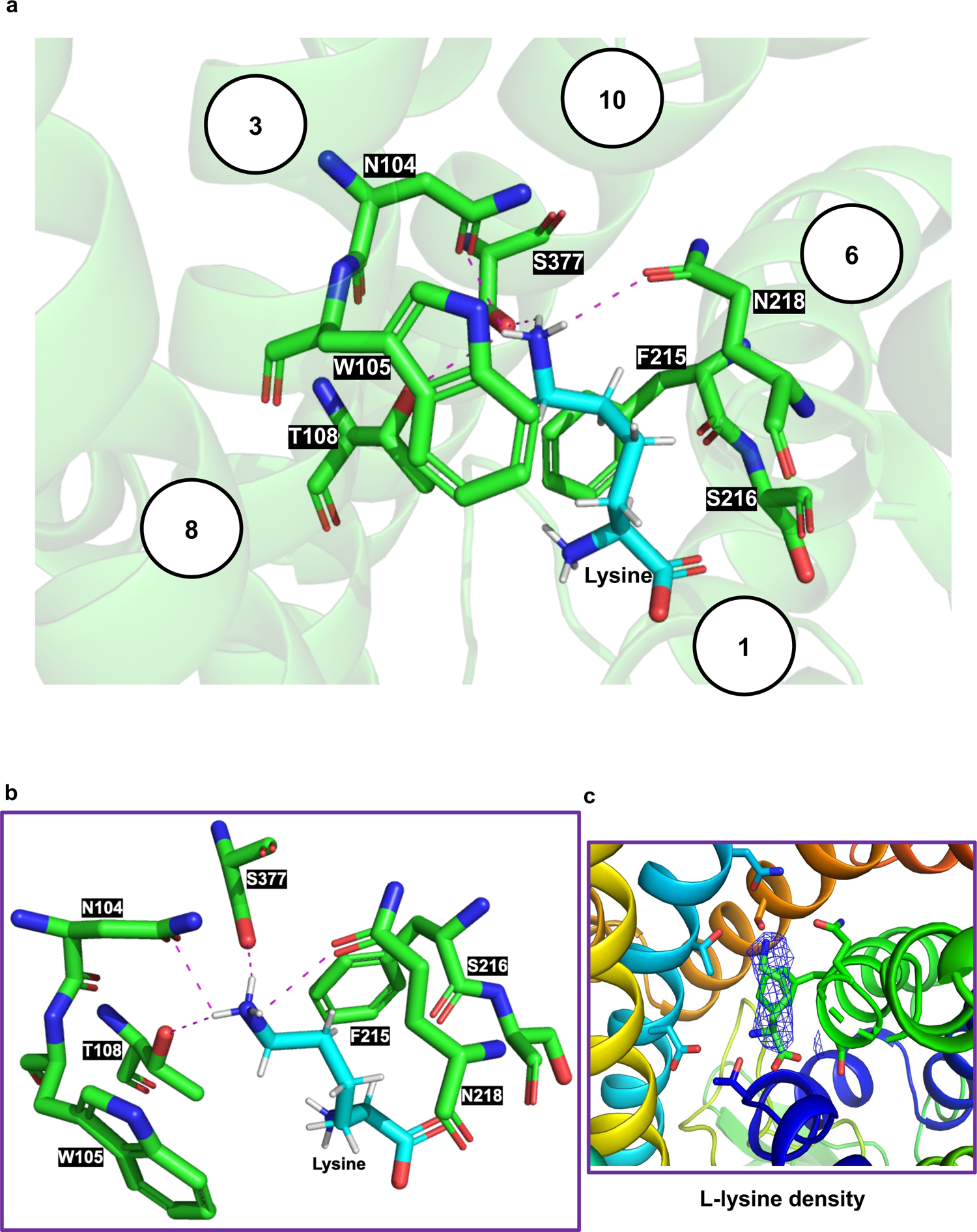
L-lysine binding site of *Pseudomonas aeruginosa* LysP. **a,** L-lysine binding site of LysP viewed from the cytoplasm. The L-lysine is coordinated to LysP by residues on TM1, TM3, TM6 and TM10 through hydrophobic interactions, a cation ρε-interaction, and by hydrogen bonding. **b**, Coordinating residues in the L-lysine binding site of LysP as shown in **a**. **c**, LysP binding site showing the L-lysine electron density map.

This cryo-EM structure displayed a 5 + 5 inverted repeat topology in which the first 5 transmembrane helices (TM1 - TM5) are a repeat of the next 5 (TM6 - TM10), with TM1 and TM6 broken in the middle^39,43^ (Fig. 3a and Extended Data Figs. 3b). TM2 is linked to TM3 by a lateral helix (LH) located in the cytosol (Fig. 3a & b and Extended Data Fig. 3b). TM10 and TM11 are linked by a β-hairpin located in the cytoplasm (Fig. 3a & b and Extended Data Fig. 3b). Unique to LysP, TM11 and TM12 are broken helices which has not been reported previously for the APC family of transporter (Extended Data Fig. 3a & b).

### Lysine binding site in LysP

The structure of the LysP-Nb5755 complex was determined in complex with L-lysine at 3.68 Å resolution by cryo-EM (Fig. 3). L-lysine is contacted by residues on TM1, TM3, TM6 and TM10 (Fig. 4a, b & c). Specifically, L-lysine is coordinated by cation-π interactions with Trp105 and by hydrophobic interactions with Phe215 (Fig. 4a & b). In addition, and unlike the other amino acid transporters, the ε-amino group of L-lysine forms hydrogen bonds with polar uncharged amino acids, that include Asn104 and Thr108 on TM3, Gln218 on TM6, and Ser377 on TM10 (Fig. 4a & b).

### Molecular basis of L-lysine selectivity by LysP

Transport and binding studies of *P. aeruginosa* LysP showed that it specifically binds and transports L-lysine (Fig. 1c & d and Extended Data Fig. 8). To understand the molecular underpinnings for the specificity of LysP towards L-lysine, sequence and structural alignments were performed with a close homologue of the mammalian CAT transporter family from *Geobacillus kaustophilus* (GkApcT)^39^ and the *Escherichia coli* arginine-agmantine antiporter from the APC transporter family (EcAdiC)^25,26^ (Fig. 5a, b & c). The structures of GkApcT^39^ and EcAdiC^25,26^ were resolved in complex with L-arginine. The residues that coordinate L-arginine in both GkApcT and EcAdiC (Extended Data Fig. 6c & d) ^39^ are different from those that coordinate L-lysine in LysP (Fig. 4a & b, Extended Data Fig. 6a); of note is the Trp105 in LysP which is homologous to Tyr116 in GkApcT and Cys97 in EcAdiC in TM3 (Fig. 5a & b). We propose that LysP utilizes Trp105 as a gate that sterically clashes with the guanidinium group of arginine during the transport cycle (Fig. 5b). By contrast, the ε-amino group of L-lysine is accommodated in the binding pocket because it is smaller than the guanidinium of arginine (Fig. 5b & d). Structural analysis on EcAdiC reported a hydrogen bond formed between the guanidinium group of L-arginine and the Cys95 on TM3 with Trp293 on TM6 instead acting as its gating residue in the occluded state^25^ (Fig. 5c and Extended Data Fig. 6c).

**Fig. 5.**
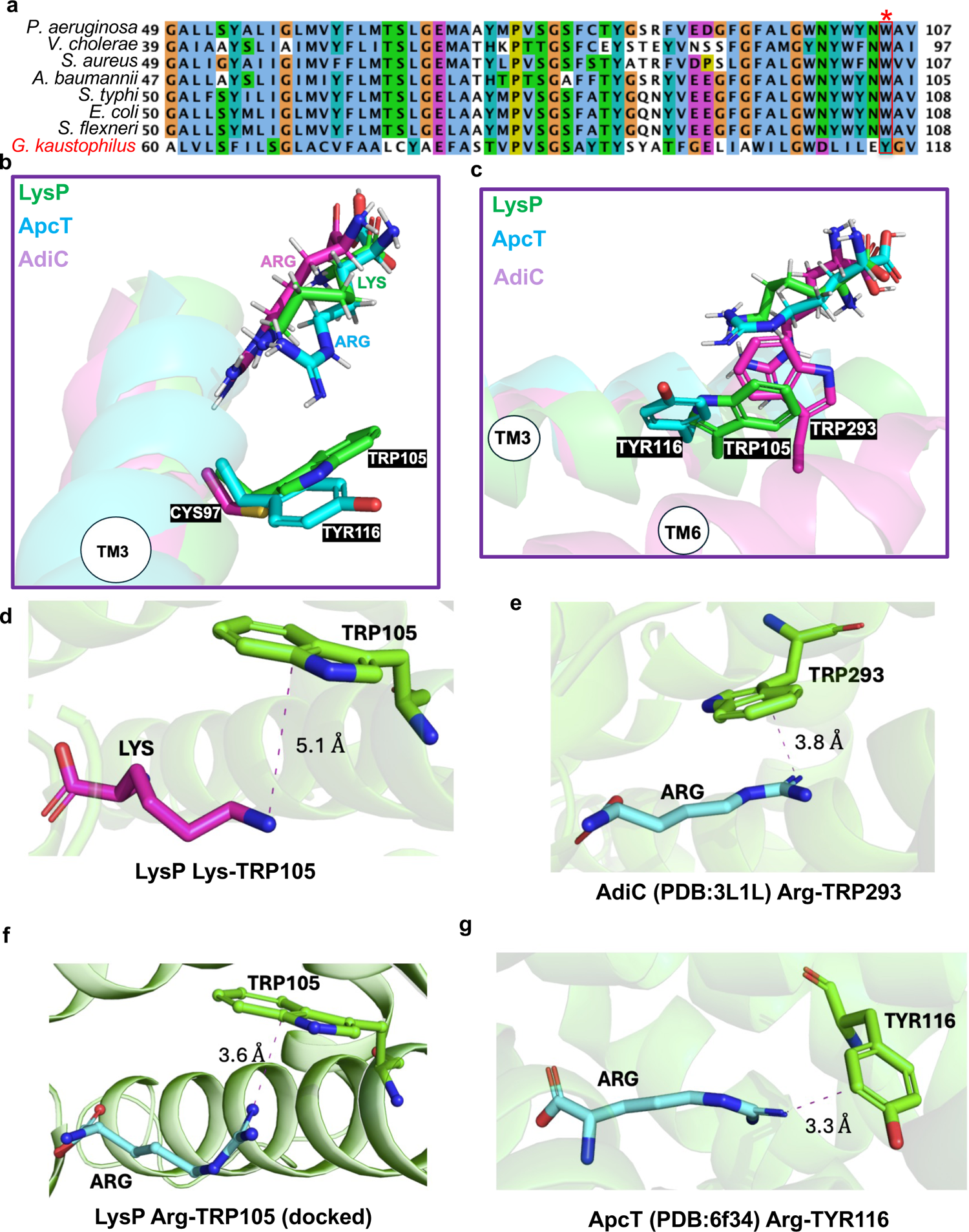
Molecular basis of L-lysine selectivity by LysP. **a,** Sequence alignment of LysP from different bacterial species and *Geobacillus kaustophilus* ApcT. W105 in LysP is homologous to Y116 in ApcT. **b,** Structural comparison of LysP (green), ApcT (cyan) and AdiC (pink) showing the cation-π interaction between L-lysine in LysP and Trp105 on TM3. LysP Trp105 is homologous to ApcT Tyr116 on TM3 which forms cation-π interaction with L-arginine. In AdiC, the corresponding residue is Cys95 on TM3 which forms a hydrogen bond with the guanidinium group of L-arginine instead. **c**, Structural comparison of LysP (green), ApcT (cyan) and AdiC (pink) highlighting the corresponding gating residue, Trp293 on AdiC located on TM6 instead of TM3 as in LysP. **d**, Cation-π interaction between the c-amino group of L-lysine and the gating residue, Trp105 in LysP. **e**, Parallel or stacked cation-π interaction between L-arginine and the gating residue, Trp293 in AdiC (PDB: 3L1L). **f**, Perpendicular or T-shaped cation-π interaction between L-arginine docked in LysP and the gating residue, Trp105. **g**, Parallel or stacked cation-π interaction between L-arginine and Tyr116 in ApcT (PDB: 6f34).

**Fig. 6.**
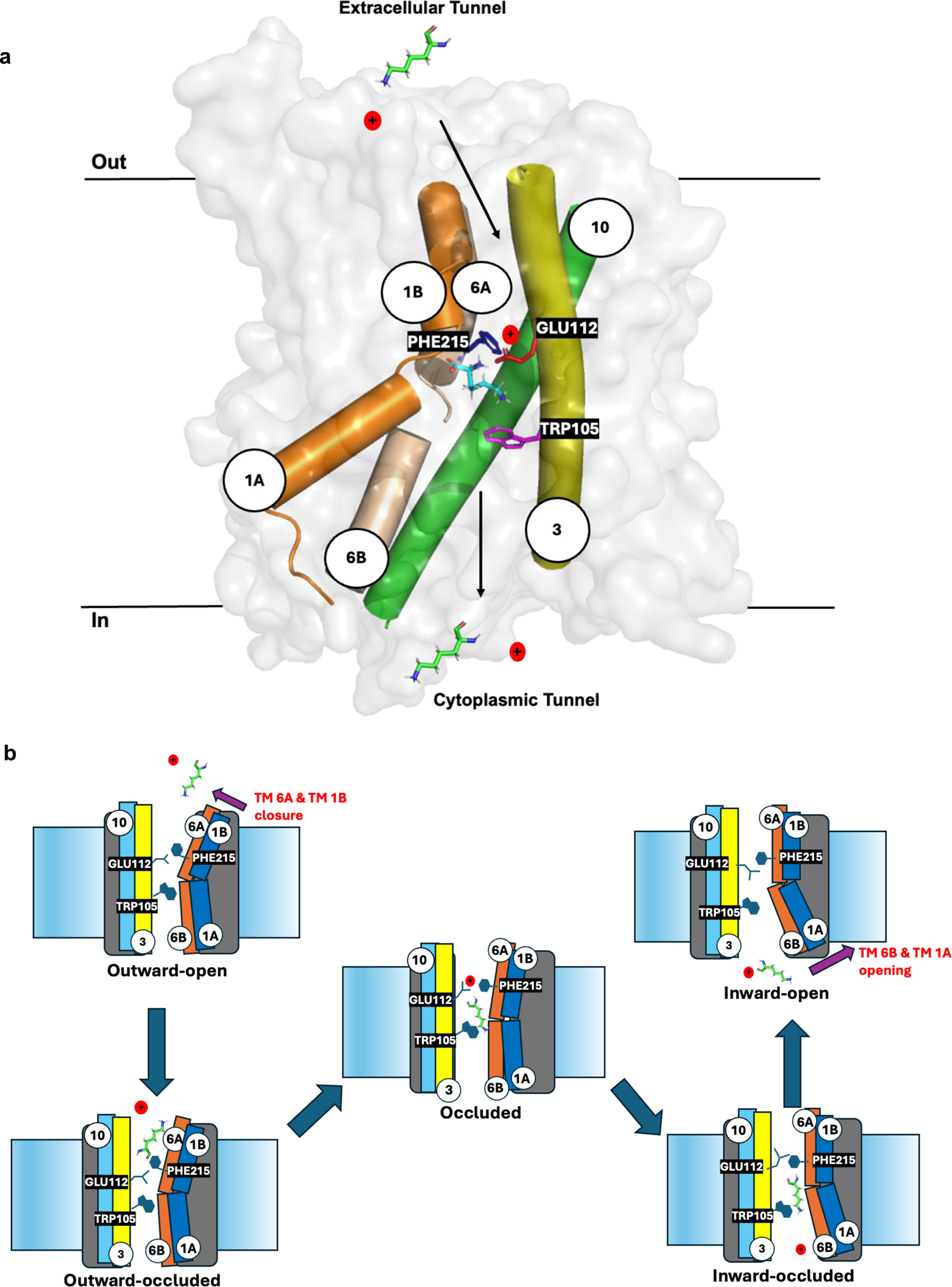
Proposed mechanism for L-lysine transport by LysP. **a**, Surface view of gating residues Trp105, Phe215 and protonated Glu112 and transmembrane helices TM1, TM3, TM6 and TM10 which form the lysine transport channel and the major gating helices. **b**, In the outward state of LysP, the extracellular tunnel is open to a proton and an L-lysine molecule, guiding them to the binding site. The binding of L-lysine and the protonation of Glu112 on TM3 result in the movement of TM6 towards TM3, forming a strong cation-π interaction between Phe215 on TM6 and the protonated Glu112 of TM3. This conformational change facilitates the closure of the extracellular gate and the formation of the occluded state. A strong cation-π interaction between the ε-amino group of the L-lysine substrate and Trp105 on TM3 follows, causing further conformational changes that drive LysP into a fully occluded state and lead to the opening of the intracellular gate. The opening of the intracellular gate is further facilitated by the release of the proton to the cytoplasm from Glu112, which causes movement of TM1 and TM6 away from the cytoplasmic tunnel, allowing the release of L-lysine and the proton.

In addition, docking studies revealed similar residues coordinate L-arginine in LysP as L-lysine (Extended Data Fig. 6b). However, the guanidinium group of L-arginine interacts with Trp105 in a more unfavorable perpendicular or T-shaped cation-π interactions (Fig. 5f). The unfavorable T-shaped interaction^44^ between L-arginine and Trp105 in LysP, unlike the more favorable parallel or stacked cation-π interactions seen between L-arginine and Trp293 in EcAdiC (Fig. 5e) and Y116 in GkApcT (Fig. 5g), could be the reason for the low binding and transport activity observed with LysP for L-arginine compared to L-lysine (Fig 1c & d and Extended Data Fig. 7c).

The specific transport of L-lysine by LysP is likely a survival mechanism. When the bacteria are exposed to extremely low pH, transport of L-lysine is activated. L-lysine, in turn, upregulates the *cadBA* operon enabling lysine decarboxylation and subsequent cadaverine export which neutralizes the extreme acidity outside the cell contributing to the survival of the bacteria (Fig. 1a). Indeed, the *cadBA* operon can only be induced in the presence of L-lysine which is shuttled into the cytoplasm by LysP^12^ and in addition, by CadC upon it’s interaction with LysP after sensing a low pH in the periplasm^13^.

Finally, the L-lysine in LysP is sandwiched between Phe215 and Trp105 (Fig. 4a & b). While the latter act as a gating residue in the inward-occluded state, we propose that the former acts also as a gating residue in the outward-open state to prevent the release of substrate back into the periplasm. In EcAdiC, Trp202 which is homologous to Phe215 in LysP, has been reported to be the gating residue in the outward-open state^25,26^. Both are located on TM6.

### LysP specifically binds and transports L-lysine using a proton gradient

To establish whether LysP is a lysine-specific transporter, LysP was reconstituted into liposomes and the transport of tritiated (^3^H) L-lysine was measured in the presence of an excess of the twenty cold proteinogenic amino acids including L-4-thialysine and D-lysine. The robust transport of ^3^H-[L-lysine] (Fig. 1b) was inhibited only by cold L-lysine and L-4-thialysine and not by the other amino acids tested, indicating that LysP is a lysine-specific transporter (Fig. 1c). The transport inhibition data performed in liposomes agrees with binding data obtained by microscale thermophoresis (MST). The *K_d_* for L- and D-lysine was 14 and 151 μM, respectively (Fig. 1d, Extended Data Fig. 7a & b) and for L-4-thialysine, 31 μM (Fig. 1e). The *K_d_* for L-arginine was 15 mM (Extended Data Fig. 7c), which explains why no binding was seen by MST for L-arginine in the concentration range utilized for L-lysine (Fig. 1d). Proteoliposomes reconstituted with a proton gradient (pH 7 in and pH 4 out) showed robust transport for L-lysine (Extended Data Fig. 7e). No significant transport was observed with proteoliposomes reconstituted either without a proton gradient (pH 4 in and pH 4 out) (Extended Data Fig. 7c) or with the addition of a sodium gradient (Extended Data Fig. 7f). These results demonstrate that LysP utilizes a proton gradient for L-lysine transport.

### LysP nanobody CA5755 binding interface

LysP loop regions on the periplamic surface of the transporter interact with the Nb5755 nanobody. The interface between the nanobody and LysP results in a buried surface area of 793 Å^2^ and 882 Å^2^ for the nanobody and LysP, respectively, which is in the range of a typical protein-protein interface^45^ (Extended Data Fig. 5a & b). The nanobody contributes ∼11 residues located on CDR1, CDR2 and CDR3 (Extended Data Fig. 5a & b), whereas LysP provides 29 residues on loops and helices at the interface (Extended Data Fig. 5b). There are several hydrophobic patches consisting of aliphatic and aromatic amino acids from both nanobody and LysP (Extended Data Fig. 5b). One of the key determinants for the stability of protein– protein complex association is the formation of a hydrophobic core or hydrophobic contacts among nonpolar amino acids^46^. This contributes to explaining why the nanobody–LysP interaction is so strong (as judged by an increase in melting temperature by 20 °C upon complex formation, Fig. 2a) (Extended Data Fig. 5b). The interface region is rich in polar contacts too. These include six hydrogen bonds and three salt bridges (Extended Data Fig. 5b). One of the important polar residues is Arg26 from the nanobody (Arg26n) making one hydrogen bond and two salt bridges with the negatively charged carboxylate group Glu288 of LysP (Extended Data Fig. 5b). Arg52n forms electrostatic interaction with Asp277 (Extended Data Fig. 5b) and hydrogen bonds with Asn269 and Asp277 in LysP (Extended Data Fig. 5b). Other hydrogen bonds include the side chains of Tyr31n, Asn76n, and backbone carbonyl oxygen of Phe28n from the nanobody and the side chain of Asn19, backbone carbonyl of Ser276 and side chain of Arg289, respectively (Extended Data Fig. 5b).

## Discussion

The structural and functional analysis of *P. aeruginosa* LysP has revealed three key molecular aspects of LysP function and pharmacological potential.

### LysP selectively transports L-lysine

The cryo-EM structure of the LysP–Nb5755 complex was resolved with bound L-lysine in an inward-occluded state (Fig. 2g, and Fig. 3a & b). This is, to the best of our knowledge, the first structural characterization of a bacterial cationic amino acid transporter that specifically recognizes and transports L-lysine. Indeed, most amino acid transporters exhibit broad spectrum specificity or substrate promiscuity. We propose that the L-lysine specificity of LysP originates principally from unfavourable interactions between the side chains of other amino acids and the gating residue, Trp105 located on TM3 of LysP but forms strong cation-π interactions with the ε-amino group of L-lysine. However, the guanidinium group of arginine is bulkier and may introduces a steric clash or form an unfavorable T-stacked interaction with this Trp105. This is evident from the *in vitro* MST study in which L-arginine binding was more than a thousand-fold weaker than that of L-lysine (Fig. 1d, Extended Data Fig. 7c). Also, in the transport inhibition study, L-arginine did not inhibit the transport of L-lysine at a concentration of 10 mM (Fig. 1c). In contrast, in the crystal structure of GkApcT, a cationic amino acid transporter in complex with L-arginine, Tyr116 (corresponding to Trp105 in LysP) interacts with the guanidinium group of arginine in a parallel or stacked type arrangement (Fig. 5b & g). Likewise, in the arginine-agmantine antiporter, EcAdiC, L-arginine interacts with the gating Trp293 residue in a more favorable stacked alignment. We also proposed that the residues coordinating the amino acid side chains may play a crucial role in determining the orientation of these side chains towards either perpendicular or parallel cation-π interactions. This, in turn, could significantly influence binding affinity and specificity. Indeed, in LysP, the ε-amino group is predominantly coordinated by polar uncharged amino acids. This coordination pattern contrasts with that observed in EcAdiC and GkApcT, as illustrated in Fig. 4a & b and Extended Data Fig. 6c & d. Finally, to support the specificity of LysP for L-lysine, transport assays revealed that no other proteinogenic amino acid was able to compete for the transport of L-lysine (Fig. 1c).

### Proposed lysine-specific uptake mechanism across bacterial cell membrane for extreme pH regulation and survival

An analysis of LysP structure suggests that this lysine transporter shares a similar alternating access transport mechanism with GkApcT^39^ and EcAdiC^24–26^. In the outward state of LysP, the extracellular tunnel is opened to a proton and L-lysine molecule and channels them to the binding site (Fig. 6a & b). The binding of L-lysine and the protonation of Glu112 on TM3 results in movement of TM6 towards TM3 to form a strong cation-π interaction between its Phe215 on TM6 and the protonated Glu112 of TM3. This conformational change facilitates closure of the extracellular gate and the formation of the occluded state (Fig. 6b), followed by a strong cation-π interaction between the ε-amino group of the L-lysine substrate and Trp105 on TM3 (Fig. 6a & b). This cation-π interaction causes further conformational changes that drive LysP into a fully occluded state and lead to the opening of the intracellular gate. Opening of the intercellular gate is further facilitated by proton release to the cytoplasm from Glu112, that causes movement of TM1 and TM6 away from the cytoplasmic tunnel for the release of L-lysine and the proton (Fig. 6a & b). Our cryo-EM structure represents the inward-occluded state where the cation-π interactions between Trp105 on TM3 and the ε-amino group of the L-lysine substrate (Fig. 6a & b) is intact. However, the cation-π interaction between Phe215 on TM6 and the protonated Glu112 on TM3 is broken due to the deprotonation of Glu112. Finally, in our structure, the TM1A and TM6B has moved away from the cytoplasmic tunnel to open the binding site ready for the release of L-lysine and proton to the cytoplasm (Fig. 6a & b). Water binding has been established to be crucial for EcAdiC stabilization, and for shaping the substrate-binding site with waters acting as placeholders for substrate atoms^47^. Likewise, it is possible that water binding plays a role in LysP transport.

### Antibiotic target potential

Previous studies have demonstrated that the addition of specific amino acids such as lysine, arginine, or glutamate can enable the recovery and growth of bacteria that have been exposed to extremely low pH conditions^43^. LysP drives lysine import and upon interaction with CadC, LysP triggers the upregulation of the *CadBA* operon leading to CadA and CadB expression^9–13^. CadA utilizes a proton to decarboxylate lysine forming cadaverine while CadB exports the alkaline product into the extracellular milieu to neutralize local acidity^9–13^. The expression and function of CadA and CadB, is dependent on the presence of L-lysine and the interaction of LysP and CadC^13^. Inhibiting lysine transport by LysP and/or its interaction with CadC would deprive the bacteria of a resistance mechanism whereby debilitating acidity is neutralized (Fig. 1a).

Armed with the structural and functional insights revealed in this study, LysP emerges as a promising target for the development of antibiotics against *P. aeruginosa* and potentially other bacterial species (Extended Data Fig. 1a & b). Inhibition of LysP could disrupt essential cellular processes required to counter extremely acidic environmental conditions that hamper bacterial growth. In this study, thialysine, an antibacterial lysine analog that targets bacterial translation^48–50^, was identified as an inhibitor of lysine transport by LysP (Fig. 1c). It works because the transporter and other systems cannot discriminate between L-lysine and L-4-thialysine. Thialysine can be incorporated into proteins^51–53^, causing them to become unstable and rapidly degraded^54^. Since thialysine competitively inhibits the transport of lysine by LysP (with a *K_i_* of approximately 31 μM), it can be further exploited to inhibit bacterial growth by targeting LysP. Indeed, the growth of *E. coli* was completely inhibited at a 5 μM concentration of thialysine^49^. This suggests that targeting bacterial LysP with thialysine and other lysine analogs could be a promising strategy for developing new antibacterial agents.

## Methods

### Cloning and overexpression of LysP

The gene encoding the lysine specific permease (*lysP*) from *P. aeruginosa* strain PAO1 was cloned into pET200/D-TOPO (Invitrogen), digested with NdeI and BamHI restriction enzymes and then sub-cloned into the standard pET 28(a) derived GFP-His_8_ fusion vector (pWaldo-GFPd)^55^ between NdeI and BamHI restriction sites. The forward and reverse primers used for the PCR amplification were 5’-CACCCATATGACTGACCTGAACACCAGCCAG-3’ and 5’-GGATCCGGTATTGGTCGGGCTGACGTC-3’ respectively. The template DNA source used for the PCR reaction was from *P. aeruginosa* strain PAO1 cells grown in Luria Bertani (LB) broth medium (Fluka). The inclusion of the *lysP* gene in the pWaldo-GFPd vector was validated by DNA sequencing, using T7 forward and reverse primers (Eurofins MWG GmbH, Anzinger Strasse 7a, D-85560 Ebersberg, Germany).

A single colony of freshly transformed *Escherichia coli* (DE3) C43 cells with pWaldo-GFPd vector harboring the *lysP* gene from *P. aeruginosa* strain PAO1 grown on LB agar/kanamycin plate was used to inoculate 50 mL of LB supplemented with 50 μg/mL of kanamycin and grown for 16 h at 37 °C with shaking at 200 r.p.m using the INFORS HT multitron standard shaker. The culture was then used to inoculate (1/50 dilution) 3 x 1 L of Terrific Broth media supplemented with 50 μg/mL of kanamycin and grown at 37 °C with shaking at 200 r.p.m using the INFORS HT multitron standard shaker. The temperature of the shaker was dropped from 37 °C to 25 °C after induction with 0.4 mM isopropylthio-β-galactoside (IPTG) at an OD_600_ of 0.6. The cells were then grown at 25 °C with shaking at 200 r.p.m for a further 22 h. GFP fluorescence (excitation 485 nm and emission 512 nm) was read on 100 μL samples taken 22 h after IPTG induction using the SpectraMax Gemini EM plate reader (Molecular Devices).

### Membrane isolation and purification of LysP

3 L of over-expressed cells were harvested immediately by centrifugation (Beckman Coulter Avanti J-20 XPI centrifuge, Beckman JLA-8.100 rotor) at 6,000 x g for 10 min at 4 °C. Supernatants were discarded and cell pellets were resuspended in ice cold 1 x phosphate buffered saline at pH 7.4 (PBS). Prior to cell lysis, 1 mM MgCl_2_, pefabloc (1 mg/mL final concentration) and DNase (20-100 U/mL final concentration) was added to the cells and the cells were broken using a Constant System cell disruptor (2 passes at 30 kPSI). The unbroken cells and debris were removed by centrifugation at 20,000 x g for 10 min at 4 °C (Beckman Coulter Avanti J20 XPI centrifuge, JLA-16,250 rotor). The supernatant containing the membrane fraction was transferred to Beckman ultracentrifuge tubes and subjected to ultracentrifugation for 2 h at 41,000 r.p.m (Beckman Optima L-100 XP ultracentrifuge, Beckman 45 Ti rotor). Membrane pellets were resuspended to a total protein concentration of 3.5 mg/mL in ice cold 1 x PBS using the Bicinchoninic Acid (BCA) protein assay kit (Pierce), rapidly frozen in liquid nitrogen and stored at −80 °C until used.

Membrane suspensions were thawed at room temperature (22 °C) and then diluted to a total protein concentration of 3.0 mg/mL in solubilization buffer (1 x PBS, 10 %(v/v) glycerol, 150 mM NaCl and 1%(w/v) of n-dodecyl-β-D-maltopyranoside (DDM)) and was further incubated for 1 h at 4 °C with gentle stirring. Non-solubilized materials were removed by centrifuging the sample at 41,000 r.p.m for 1 h at 4 °C using rotor type 45 Ti in the OptimaTM L-100 XP Ultracentrifuge (BECKMAN COULTER). The resulting supernatant containing the solubilized LysP-GFP was incubated with Ni^2+^-NTA resin (Qiagen) for 2 h at 4 °C using 1 mL of resin per mg GFP. To reduce non-specific binding, 10 mM imidazole pH 7.5 was added to the sample. The slurry was loaded on to a glass Econo gravity column (Bio-Rad) and washed with 20 column volumes of solubilization buffer containing 10 mM imidazole pH 7.5. The column was washed with a further 20 column volumes each of solubilization buffer containing 20 and 35 mM imidazole pH 7.5 at 4 °C. The protein was then eluted from the column and the eluate collected using 40 mL elution buffer (solubilization buffer containing 250 mM imidazole pH 7.5) at 4 °C. The eluate was incubated overnight at 4 °C with an equal amount of His-tagged Tobacco etch virus (TEV) protease (1 mg TEV protease/mg GFP) was added into a dialysis tube (14 kDa MWCO) and the sample was dialysed against 3 L of dialysis buffer containing 20 mM Tris HCl pH 7.5, 150 mM NaCl and 0.003 %(w/v) lauryl maltose neopentyl glycol (LMNG) at 4 °C with gentle stirring of the dialysis buffer with a magnetic rod/stirrer. The sample consisting of GFP-free LysP, GFP, TEV and likely a small amount of uncleaved LysP-GFP was filtered using 0.22 μm Millipore filters to remove protein aggregates and was further subjected to the second round of IMAC to remove the free GFP, TEV protease and any uncleaved LysP-GFP. In this reverse IMAC step, a 5 mL Ni^2+^-NTA HisTrap fast flow column (Qiagen) was equilibrated with 10 mL of buffer containing 20 mM Tris HCl pH 7.5, 150 mM NaCl and 0.003 %(w/v) LMNG at a flow rate of 0.2 mL/min at 4 °C and the eluent containing GFP-free LysP was collected and concentrated at 3,500 x g at 4 °C using rotor C0650 in the AllegraTM X-22R centrifuge (BECKMANN COULTER Clare, Ireland) for 5 min at 4 °C using a Centricon 50 kDa molecular weight cut off (MWCO) concentrator (Millipore: MA, USA, Lot R0KA67379). After the 5 min spin, the sample was resuspended by pipetting up and down. The concentrated sample (0.5 mL final volume) was loaded onto a Superdex 200 increase 10/300 size exclusion column (GE Healthcare) at a flow rate of 0.4 mL/min and the absorbance at 280 nm was monitored. Prior to loading the sample, the column was pre-equilibrated with buffer containing 10 mM Tris HCl pH 7.5, 100 mM NaCl and 0.003 %(w/v) LMNG. For transport assays, the 0.003%(w/v) LMNG in the gel filtration buffer was substituted with 0.03%(w/v) DDM. LysP was concentrated using 100 kDa molecular weight cut off concentrator (Millipore). The concentration of purified GFP-free LysP was determined from the absorbance at 280 nm, using an extinction coefficient of 97,290 M^-1^.cm^-1^ calculated from the amino acid sequence.

### Circular dichroism

CD spectra of purified LysP at 0.6 mg/mL in 150 mM NaCl, 20 mM Tris pH 7.5 and 0.003 %(w/v) LMNG, was recorded using a Jasco J-815 CD Spectrophotometer from 190 nm to 260 nm at a scan speed of 100 nm/min, a band width of 1 nm, a path length of 1 mm and an ellipticity of 0.1 degrees at 20 °C. The same parameters were used for the blank containing 150 mM NaCl, 20 mM Tris pH 7.5 and 0.003 %(w/v) LMNG without LysP. Data were processed using the K2d method in the online CD analysis webpage (DIOCROWEB) (<u>htt://dichroweb.cryst.bbk.ac.uk/html/home.shtml</u>)^56–58^.

### Microscale thermophoresis (MST)

MST was carried out as described previously^59,60^. This experiment was carried out at NanoTemper Technologies GmbH, Floβergasse 4, 81369 Munich, Germany. In brief, serial 2-fold dilutions of each ligand were prepared starting at 800 µM for L-lysine, 6.4 mM for L-4-thialysine, 50 mM D-lysine, 200 mM of L-arginine and 100 mM each of L-histidine, glycine, L-proline, L-glutamine, L-glutamate, L-methionine, L-isoleucine, L-leucine, L-threonine, L-valine and L-alanine to obtain 16 different concentrations of ligand at 22 °C. All concentrations of ligands were mixed with protein in a 1:1 by volume ratio and 4 µL each were loaded into glass capillaries. The final protein concentration was 0.6 µM. MST analysis was carried out using Monolith Label Free instrument (NanoTemper) using a laser power and LED sensitivity of 20 % at 22 °C. All ligands and protein solutions used for MST were prepared in buffer containing 150 mM NaCl, 20 mM Tris pH 7.5 and 0.003 %(w/v) LMNG.

### Reconstitution and counterflow transport assays

The reconstitution and counterflow assay begin with the purification of the lipid to be used for reconstitution with acetone and ether. 100 mg of *E. coli* polar lipid (Avanti) (white solid powder) was dissolved in 2 mL of chloroform and mixed with 10 mL of acetone under argon containing 2 μL of β-mercaptoethanol (14.7 M stock) and the mixture was stirred for 16 h at 4 °C. The precipitated lipid was sedimented by centrifuging at 10, 000 x g for 12 min at room temperature and the supernatant discarded. The remaining lipid pellet was resuspended in 10 mL of diethyl-ether under argon containing 2 μL of β-mercaptoethanol (14.7 M stock) and centrifuged at 8,000 x g for 10 min at room temperature. The supernatant containing the dissolved lipid was retained and the ether removed by evaporation using argon, to produce a lipid film. The purified lipid was resuspended in chloroform at a final concentration of 10 mg/mL, under argon and stored at −20 °C. Glass tubes were used throughout at room temperature as chloroform dissolves plastic tubes. The purified lipid extract stored in chloroform was dried under argon and the lipid film was rehydrated to a final concentration of 10 mg/mL using reconstitution buffer (50 mM potassium phosphate pH 7.6, 20 mM cold L-lysine, 1 mM DTT) by a sequential series (10 times) of vortexing (30 s) and heating at 50 °C. To aid rehydration a series of incubations at 50 °C and mixing were performed until the lipid film had completely dispersed. Unilamellar vesicles of size, 0.1 μm were prepared using an extruder (Lipex Biomembranes, Inc. Vancouver, British Columbia) at a pressure of 300 psi to pass the lipid solution once through 0.4 μm polycarbonate Nucleopore^TM^ Track Etch double membrane filters, it was then passed once through 0.2 μm polycarbonate Nucleopore^TM^ Track Etch double membrane filters and 7 times through 0.1 μm pore size polycarbonate Nucleopore^TM^ Track Etch single membrane filters. The extrusion was carried out with the extruder kept at 50 °C in a water bath. This produced a homogenous suspension of 0.1 μm diameter vesicles. The liposomes were used immediately for reconstitution of LysP. For reconstitution, β-octyl-glucoside (OG) was used to destabilise the preformed liposomes and to allow the insertion of LysP into the liposome membrane. 1 mL of lipid at 10 mg/mL in reconstitution buffer was mixed at 4 °C for 15 min with 1.25 %(w/v) solution of β-OG (10 %(w/v) stock solution) and purified LysP at 10 mg/mL in 150 mM NaCl, 20 mM Tris-HCl pH 7.5 and 0.03 %(w/v) DDM (100:1 lipid to LysP weight ratio) in a 5 mL glass test tube. For the control (liposomes alone), an equal volume of Buffer A (10 μL) was added in place of purified LysP (10 mg/mL in 150 mM NaCl, 20 mM Tris-HCl pH 7.5 and 0.03 %(w/v) DDM). Excess β-OG was removed from the mixture by diluting with 70 mL of reconstitution buffer and the sample was centrifuged for 1 h at 100,000 x g at 4 °C to pellet the proteoliposomes. The supernatant was discarded carefully, the proteoliposome pellets were resuspended in 15 μL reconstitution buffer on ice and used directly.

For counterflow assay, proteoliposomes or liposomes were diluted 1 in 25 (by vol) using counterflow buffer (CB, 50 mM potassium phosphate pH 7.6 and 1 mM DTT) and the counterflow assay^61^ was initiated by adding 40 µL of [^3^H] L-lysine (stock, 50 µM, 100 µCi/mL). After the addition of [^3^H] L-lysine, samples (80 µL) were taken at different time points and applied to a 0.22 µm pore size MF-Millipore™ Membrane Filter. The filters were immediately washed under vacuum with 4 mL of ice-cold CB. The washed filter was transferred to a 20 mL scintillation vial and submerged in 10 mL emulsifier safe liquid scintillant. The level of radioactivity associated wtih the liposomes or proteoliposomes was measured by liquid scintillation counting. All protein batches used for reconstitution were purified in DDM. For Na^+^ gradient tests, 50 mM NaCl was added to the external CB with the internal CB kept constant (50 mM potassium phosphate pH 7.6 and 1 mM DDT), while for H^+^ gradient tests, the external pH was kept constant at pH 4 using 25 mM citrate-phosphate buffer while the internal pH used were pH 4 and pH 7.0 using 25 mM citrate-phosphate buffer instead of 50 mM KPi pH 7.6. An inhibition study was carried out at external pH 4.0 and internal pH 7.0 in the presence of 10 mM external cold inhibitors (the twenty L-amino acids, L-4-thialysine and D-lysine) and samples were collected and analysed after 3 and 20 min.

### Electron microscopy sample preparation and data acquisition

The cryo-EM grids were prepared by applying 3 μL of the LysP-Nb5755 complex protein at 2 mg/mL in buffer containing 100 mM NaCl, 20 mM Tris-HCl pH 7.5, 0.03 %(w/v) DDM and 10 mM L-lysine, to a glow-discharged Quantifoil R1.2/1.3 200/300-mesh copper/gold holey carbon grid (QUANTIFOIL, Micro Tools GmbH, Germany) and blotted for 3.0 s under conditions of 100% relative humidity and 4°C before being plunged into liquid ethane using a Mark IV Vitrobot (FEI). Grids were screened in a 200 keV Talos Arctica microscope (Thermo Fisher) at Uppsala University, Sweden and the Membrane Protein Lab, Diamond Light Source Ltd., UK cryo-EM imaging facilities. Micrographs were acquired on a Titan Krios microscope (FEI) operated at 300 kV with a K2 Summit direct electron detector (Gatan) at the cryo-EM Swedish National Facility in Stockholm. SerialEM software was used for automated data collection following standard procedures^62^. A calibrated magnification of ×130,000 was used for imaging, yielding a pixel size of 0.67 Å on images. The defocus range was set from −0.8 to −2.4 μm. Each micrograph was dose-fractionated to 50 frames under a dose rate of 7 - 8 electrons.pixel^-1^.s^-1^, with a total exposure time of 8 s, resulting in a total dose of ∼80 electrons.Å^-2^.

### Cryo-EM data processing, model building, and structure analysis

The data set was processed using CryoSPARC^63^. The dose fractionated movie frames were aligned using “patch motion correction”, contrast transfer function (CTF) were estimated using “Patch CTF estimation”, the particles were picked using automated blob picker and extracted by binning 6 times in cryoSPARC live. Afterwards, several rounds of 2D classifications were performed. Finally, good particles were selected and extracted by binning equal to 1.0 Å.pixiel^-1^. 177,424 particles were further used for *ab initio* maps and heterogeneous refinements. After refinement, 145,063 particles were selected. Several rounds of non-uniform refinement and local refinement were performed. To remove bad particles, further phase randomized heterogeneous refinement was performed with low-pass filter 8 Å, 20 Å and 40 Å. The 78,459 particles were selected and local refinement with a tight mask was performed. The resolution reached 3.68 Å at the gold standard FSC resolution value of 0.143. The local resolution was calculated in CryoSPARC^63^. An AlphaFold 2^64^ model of LysP was automatically fitted into the cryo-EM density map of our inward-facing state. Iterative model building and real space refinement were performed using COOT^65^ and PHENIX.refine^66^. The refinement statistics are summarized in Table 1. For generating structural figures PyMOL version 3.0.1^67^ and UCSF chimera^68^ or chimeraX^69^ were used.

### Nanobody generation, expression and purification

Nanobodies (Nb5753, Nb5755, Nb5758, Nb5760, Nb5761, Nb5764 and Nb5765) against LysP were generated using previously published protocols^70^. In brief, one llama (Lama glama) was six times immunized with LysP (1 mg/mL in DDM purification buffer). Four days after the final boost, blood was taken from the llama to isolate peripheral blood lymphocytes. RNA was purified from these lymphocytes and reverse transcribed by PCR to obtain the cDNA of the open reading frames coding for the nanobodies. The resulting library was cloned into the phage display vector pMESy4 bearing a C-terminal His_6_ tag and a CaptureSelect sequence tag (Glu-Pro-Glu-Ala). Nanobodies were selected by biopanning. For this, LysP in detergent solution was solid phase coated directly on plates. LysP specific phage were recovered by limited trypsinization, and after two rounds of selection, periplasmic extracts were made and analysed using ELISA screens. Nanobodies were expressed in *E. coli* Bl21 (DE3) for subsequent purification from the bacterial periplasm. After Ni^2+^-NTA (Qiagen) affinity purification, nanobodies were further purified by size exclusion chromatography in buffer containing 20 mM Tris-HCl pH 7.5 and 150 mM NaCl. The purity of the purified nanobodies were analyse on SDS PAGE. The nanobodies were stored in the cold (4 °C) until use.

### Green fluorescent protein-thermal shift (GFP-TS) assay

The GFP-TS assay was performed as previously described^71,72^. Briefly, membranes (3.5 mg/mL in 150 mM NaCl, 20 mM Tris-HCL pH 7.5 and 1 %(w/v) DDM) were solubilized with or without 1 μM nanobody Nb5755 for 1 h at 4 °C with mild agitation. After solubilization, 1 %(w/v) β-OG was added to both samples and 120 μL aliquots of the solubilized membranes were incubated in 200 μL PCR tubes for 10 min at 4, 20, 30, 35, 40, 45, 50, 65, 70, or 100 °C using a T100^TM^ Thermal Cycler (BIO-RAD) without mixing. The heated samples were then centrifuged at 18,000 x g for 30 min at 4 °C using a Microfuge® 18 Centrifuge, Beckman Coulter. 90 μL aliquots of the supernatant were transferred into a black clear bottom 96-well plate (Nunc) and the GFP fluorescence (excitation 485 nm, emission 512 nm) measured using a SpectraMax Germini EM microplate reader (Molecular Devices).

### Molecular docking

Modeling studies were performed with the programs of the Schrödinger Small-Molecule Drug Discovery Suite 2021-4 (Schrödinger LLC, NY, USA). The 2D structures of L-lysine and L-arginine were converted to geometry optimized 3D structures of their predominant ionization states (at pH 7.4) using LigPrep (Schrödinger LLC, NY, USA). Next, the cryo-EM structure of LysP in complex with L-Lys was prepared with the Protein PrepWizard^73^. This involves adding missing hydrogen atoms, adjusting the ionization state of polar amino acids at neutral pH, and meanwhile adjusting bond orders and formal charges of the ligand, optimizing the H-bond network of the protein–ligand complex, and, finally, energetically minimizing the complex using the Optimized Potentials for Liquid Simulations (OPLS)-4 force field^74^. The geometric center of the bound L-lysine molecule was considered as the grid centroid, and flexible ligand docking was carried out using Glide with the SP scoring function^75^.

### Preparation of Figures

Graph preparation and statistical analysis were performed in GraphPad Prism 10 (for macOS). Sequence alignments were generated using Clustal Omega online software^76^. Figures with models and EM density maps were prepared in PyMOL (the PyMOL Molecular Graphics System, Version 3.0.1, Schrödinger) ^67^, UCSF Chimera^68^ and ChimeraX^69^.

### Reporting summary

Further information on research design is available in the Nature Research Reporting Summary linked to this article.

## Data availability

The structural model of the LysP-CA5755-L-lysine complex has been deposited in the Protein Data Bank under accession code 9EYD. The cryo-EM maps were deposited in the Electron Microscopy Data Bank (EMDB) under accession number EMD-50053. Materials are available upon reasonable request. Source data are provided with this paper.

---

**Extended Data Fig. 1.** Amino acid sequence alignment of *Pseudomonas aeruginosa* LysP and LysP from other bacteria orthologs. **a,** Sequence alignment of *P. aeruginosa* LysP and other enteric pathogenic bacteria (*Vibrio cholerae, Staphylococcus aureus, Acinobacter baumannii, Salmonella typhi, Escherichia coli*, and *Shigella flexneri*). **b,** Number of matching residues (percentatage identity) of the alignmnet shown in **a**.

**Extended Data Fig. 2.** Cryo-EM data processing workflow **a**, Representative EM micrograph. 10,427 micrographs were collected, and 7,297,090 particles were picked by blob picking. **b**, The selected 2D class-averages. **c**, 3D classes from heterogenous refinement and selected 145,063 particles for further processing. **d**, The first non-uniform refinement 3D map that reached 3.74 Å at the gold standard FSC value of 0.143. **e**, Heterogeneous refinement with different low-pass filtered map (20 Å or 40 Å**). f**, The local refinement using tight mask without micelle region. The final resolution reached to 3.68 Å at the gold standard FSC. **g**, Colored map according to local resolution estimation.

**Extended Data Fig. 3. a**, Cryo-EM density corresponding to representative individual transmembrane helices of LysP and L-lysine at σ = 7.0. **b**, Topology of the cryo-EM structure of LysP. LysP has 12 transmembrane helices with both C- and N-termini located in the cytoplasm. Helices 1, 6, 11 and 12 are broken in the middle, Helices 2 and 3 are linked by a lateral cytoplasmic helix and 10 and 11 by a β-hairpin.

**Extended Data Fig. 4.** Purification and screening of nanobody binders against LysP without L-lysine. **a,** Size-exclusion chromatogram of the purified nanobodies (Nb5753, Nb5755, Nb5758, Nb5760, Nb5761, Nb5764, and Nb5765). V_o_ denotes the void volume and V_t_ the included volume. n=1 for an independent experiment. **Insert,** SDS PAGE analysis of purified nanobodies. Mr = molecular weight marker. **b**, Size-exclusion chromatogram of the nanobody-LysP complex. Nanobodies were mixed with LysP in a molar ratio of 1 LysP to 3 of the respective nanobodies and incubated for 2 h on ice. n=1 for an independent experiment. **Insert,** SDS PAGE analysis of the size-exclusion chromatogram peaks. Mr = molecular weight marker.

**Extended Data Fig. 5.** LysP-Nb5755 interaction with L-lysine. **a,** Membrane view showing the LysP-Nb5755 binding interface. **b,** Enlargement of the LysP-Nb5755 binding interface showing residues in LysP and their corresponding interactions with residues on the CDR1, CDR2 and CDR3 of Nb5755.

**Extended Data Fig. 6.** Comparison of the binding site residues in LysP, *E. coli* AdiC and G. *kaustophilus* ApcT. **a,** LigPlot showing the residues coordinating L-lysine in LysP. **b,** LigPlot showing the residues coordinating L-arginine docked into the LysP structure. **c,** LigPlot showing the residues coordinating L-arginine in *E. coli* AdiC. **d,** LigPlot showing the residues coordinating L-arginine in *G. kaustophilus* ApcT.

**Extended Data Fig. 7.** Functional characterization of LysP **a,** Microscale thermophoresis (MST) binding profile of LysP to L-lysine with a *K_d_* of 14.5 μM. **b**, MST binding profile of LysP to D-lysine with a *K_d_* of 151 μM. **c**, MST binding profile of LysP to L-arginine with a *K_d_* of 15.6 mM. **d**, MST binding profile of LysP to L-histidine with a *K_d_* of 4.5 mM. **e,** pH gradient test: uptake of ^3^[H]L-lysine in reconstituted liposomes with LysP (proteoliposomes) and without LysP (control liposomes) at pH 4 in and out (red) and at pH 7 in and pH 4 out (blue). The mean ± s.e.m. of the fit is the average from four independent experiments. **f**, Sodium gradient test: uptake of ^3^[H]L-lysine in reconstituted liposomes with LysP (proteoliposome) with (blue) and without the addition of 50 mM NaCl (red). The mean ± s.e.m. of the fit is the average from four independent experiments.

**Extended Data Fig. 8.** Microscale thermophoresis (MST) analysis of LysP. **a**, MST binding profile of LysP to L-alanine, no binding was seen. **b**, MST binding profile of LysP to glycine, no binding was seen, **c**, MST binding profile of LysP to L-isoleucine, no binding was seen. **d**, MST binding profile of LysP to L-methionine, no binding was seen. **e**, MST binding profile of LysP to L-proline, no binding was seen. **f**, MST binding profile of LysP to L-valine no binding was seen. **g**, MST binding profile of LysP to L-glutamine, no binding was seen. **h**, MST binding profile of LysP to L-glutamate, no binding was seen. **i**, MST binding profile of LysP to L-threonine, no binding was seen. and **j**, MST binding profile of LysP to L-Leucine, no binding was seen. The mean ± s.e.m. of the fit is the average from two to three independent experiments.

## Acknowledgments

This research was funded by the Wellcome Trust (grant 222999/Z/21/Z to E.N.), the Wenner-Gren Foundations (grant GFOv2022-0003 to J.J.G.), the Swedish Research Council (grant 2022-02985 to J.J.G.) and the Science Foundation Ireland (grant 12/IA/1255 and 22/FFP-A/10278 to M.C.). We acknowledge the use of the Cryo-EM Uppsala facility for grid preparation and screening, funded by the Department of Cell and Molecular Biology, the Disciplinary Domains of Science and Technology and of Medicine and Pharmacy at Uppsala University. We also acknowledge the Membrane Protein Laboratory at Diamond Light Source funded by Wellcome (223727/Z/21/Z). Diamond Light Source and the Research Complex at Harwell are both Instruct-ERIC centres. Cryo-EM data was collected at the Cryo-EM Swedish National Facility funded by the Knut and Alice Wallenberg, Family Erling Persson and Kempe Foundations, SciLifeLab, Stockholm University and Umeå University. We thank Daniel Larsson, and Marta Carroni for assistance with cryo-EM data collection. We also acknowledge the support and the use of resources of Instruct-ERIC, part of the European Strategy Forum on Research Infrastructures (ESFRI), and the Research Foundation - Flanders (FWO) for their support to the Nanobody discovery. We thank Nele Buys for the technical assistance during Nanobody discovery.

## Contributions

E.N. designed and supervised the project. E.N. performed protein expression, purification, functional assays and cryo-EM sample preparation and data collection, as well as atomic model building and refinement. D.B. assisted with protein production and cryo-EM sample preparation, A.F.A.M. performed docking studies, H.C assisted with cryo-EM grid preparation, P.S. assisted with radioactivity assays. R.M. provided assistance with cryo-EM data processing. M.C. supervised the initial protein production, purification and biophysical characterization. P.J.F supervised radioactivity assays respectively. E.P. and J.S. performed nanobody generation. E.N. wrote the manuscript. All authors read and commented on the manuscript.

## Corresponding author

Correspondence to Emmanuel Nji.

## References

1. Ghebreyesus, T. A. Making AMR history: a call to action. Global Health Action 12, 1638144 (2019).

2. Lewis, K. The Science of Antibiotic Discovery. Cell 181, 29–45 (2020).

3. Nkansa-Gyamfi, N. A., Kazibwe, J., Traore, D. A. K. & Nji, E. Prevalence of multidrug-, extensive drug-, and pandrug-resistant commensal *Escherichia coli* isolated from healthy humans in community settings in low- and middle-income countries: a systematic review and meta-analysis. Glob Health Action 12, 1815272 (2019).

4. Nji, E. et al. High prevalence of antibiotic resistance in commensal *Escherichia coli* from healthy human sources in community settings. Sci Rep 11, 3372 (2021).

5. The European Centre for Disease Prevention and Control: The bacterial challenge: Time to react. Stockholm 2009.

6. European Centre for Disease Prevention and Control. Assessing the Health Burden of Infections with Antibiotic-Resistant Bacteria in the EU/EEA, 2016-2020. Stockholm: ECDC; 2022.

7. Centers for Disease Control and Prevention (U.S.). Antibiotic Resistance Threats in the United States, 2019. https://stacks.cdc.gov/view/cdc/82532 (2019) doi:10.15620/cdc:82532.

8. de Kraker, M. E. A., Stewardson, A. J. & Harbarth, S. Will 10 Million People Die a Year due to Antimicrobial Resistance by 2050? PLoS Med 13, e1002184 (2016).

9. Neely, M. N., Dell, C. L. & Olson, E. R. Roles of LysP and CadC in mediating the lysine requirement for acid induction of the *Escherichia coli* cad operon. J. Bacteriol. 176, 3278–3285 (1994).

10. Tetsch, L., Koller, C., Haneburger, I. & Jung, K. The membrane-integrated transcriptional activator CadC of *Escherichia coli* senses lysine indirectly via the interaction with the lysine permease LysP. Mol. Microbiol. 67, 570–583 (2008).

11. Rauschmeier, M., Schüppel, V., Tetsch, L. & Jung, K. New insights into the interplay between the lysine transporter LysP and the pH sensor CadC in *Escherichia coli*. J. Mol. Biol. 426, 215–229 (2014).

12. Soksawatmaekhin, W., Kuraishi, A., Sakata, K., Kashiwagi, K. & Igarashi, K. Excretion and uptake of cadaverine by CadB and its physiological functions in *Escherichia coli*. Molecular Microbiology 51, 1401–1412 (2004).

13. Küper, C. & Jung, K. CadC-Mediated Activation of the *cadBA* Promoter in *Escherichia coli*. Microb Physiol 10, 26–39 (2005).

14. Lolkema, J. S., Poolman, B. & Konings, W. N. Bacterial solute uptake and efflux systems. Current Opinion in Microbiology 1, 248–253 (1998).

15. Lim, S. S. et al. A comparative risk assessment of burden of disease and injury attributable to 67 risk factors and risk factor clusters in 21 regions, 1990–2010: a systematic analysis for the Global Burden of Disease Study 2010. The Lancet 380, 2224–2260 (2012).

16. Tacconelli, E. et al. Discovery, research, and development of new antibiotics: the WHO priority list of antibiotic-resistant bacteria and tuberculosis. The Lancet Infectious Diseases 18, 318–327 (2018).

17. Koch, C. Early infection and progression of cystic fibrosis lung disease. Pediatric Pulmonology 34, 232–236 (2002).

18. Harbarth, S. Epidemiology and Prognostic Determinants of Bloodstream Infections in Surgical Intensive Care. Arch Surg 137, 1353 (2002).

19. Osmon, S., Ward, S., Fraser, V. J. & Kollef, M. H. Hospital Mortality for Patients With Bacteremia Due to Staphylococcus aureus or Pseudomonas aeruginosa. Chest 125, 607–616 (2004).

20. Pang, Z., Raudonis, R., Glick, B. R., Lin, T.-J. & Cheng, Z. Antibiotic resistance in *Pseudomonas aeruginosa*: mechanisms and alternative therapeutic strategies. Biotechnology Advances 37, 177–192 (2019).

21. Jack, D. L., Paulsen, I. T. & Saier, M. H. The amino acid/polyamine/organocation (APC) superfamily of transporters specific for amino acids, polyamines and organocations. *Microbiology (Reading*, Engl.) 146 **( Pt** **8**), 1797–1814 (2000).

22. Shaffer, P. L., Goehring, A., Shankaranarayanan, A. & Gouaux, E. Structure and mechanism of a Na+-independent amino acid transporter. Science 325, 1010–1014 (2009).

23. Shaffer, P. L., Goehring, A., Shankaranarayanan, A. & Gouaux, E. Structure and Mechanism of a Na^+^ - Independent Amino Acid Transporter. Science 325, 1010–1014 (2009).

24. Ilgü, H. et al. Insights into the molecular basis for substrate binding and specificity of the wild-type L-arginine/agmatine antiporter AdiC. Proc. Natl. Acad. Sci. U.S.A. 113, 10358–10363 (2016).

25. Gao, X. et al. Mechanism of substrate recognition and transport by an amino acid antiporter. Nature 463, 828–832 (2010).

26. Gao, X. et al. Structure and Mechanism of an Amino Acid Antiporter. Science 324, 1565–1568 (2009).

27. Ma, D. et al. Structure and mechanism of a glutamate–GABA antiporter. Nature 483, 632–636 (2012).

28. Ma, J. et al. Structural basis for substrate binding and specificity of a sodium–alanine symporter AgcS. Proc. Natl. Acad. Sci. U.S.A. 116, 2086–2090 (2019).

29. Errasti-Murugarren, E. et al. L amino acid transporter structure and molecular bases for the asymmetry of substrate interaction. Nat Commun 10, 1807 (2019).

30. Fan, M., Zhang, J., Lee, C.-L., Zhang, J. & Feng, L. Structure and thiazide inhibition mechanism of the human Na–Cl cotransporter. Nature 614, 788–793 (2023).

31. Chew, T. A. et al. Structure and mechanism of the cation–chloride cotransporter NKCC1. Nature 572, 488–492 (2019).

32. Liu, S. et al. Cryo-EM structures of the human cation-chloride cotransporter KCC1. Science 366, 505–508 (2019).

33. Chi, G. et al. Phospho-regulation, nucleotide binding and ion access control in potassium-chloride cotransporters. EMBO J 40, (2021).

34. Xie, Y. et al. Structures and an activation mechanism of human potassium-chloride cotransporters. Sci. Adv. 6, eabc5883 (2020).

35. Yang, X., Wang, Q. & Cao, E. Structure of the human cation–chloride cotransporter NKCC1 determined by single-particle electron cryo-microscopy. Nat Commun 11, 1016 (2020).

36. Zhao, Y. et al. Structure of the human cation–chloride cotransport KCC1 in an outward-open state. Proc. Natl. Acad. Sci. U.S.A. 119, e2109083119 (2022).

37. Reid, M. S., Kern, D. M. & Brohawn, S. G. Cryo-EM structure of the potassium-chloride cotransporter KCC4 in lipid nanodiscs. eLife 9, e52505 (2020).

38. Tascón, I. et al. Structural basis of proton-coupled potassium transport in the KUP family. Nat Commun 11, 626 (2020).

39. Jungnickel, K. E. J., Parker, J. L. & Newstead, S. Structural basis for amino acid transport by the CAT family of SLC7 transporters. Nat Commun 9, 550 (2018).

40. Nji, E. Structural and Functional Studies of Pseudomonas aeruginosa Strain PAOI Lysine Specific Permease (LysP). (Trinity College (Dublin, Ireland). School of Biochemistry and Immunology, 2013).

41. Nji, E., Li, D., Doyle, D. A. & Caffrey, M. Cloning, expression, purification, crystallization and preliminary X-ray diffraction of a lysine-specific permease from *Pseudomonas aeruginosa*. Acta Crystallogr F Struct Biol Commun 70, 1362–1367 (2014).

42. Ellis, J. et al. Topological analysis of the lysine-specific permease of *Escherichia coli*. Microbiology (Reading) 141 **( Pt** **8**), 1927–1935 (1995).

43. Iyer, R., Williams, C. & Miller, C. Arginine-Agmatine Antiporter in Extreme Acid Resistance in *Escherichia coli*. J Bacteriol 185, 6556–6561 (2003).

44. Infield, D. T. et al. Cation-π Interactions and their Functional Roles in Membrane Proteins. J Mol Biol 433, 167035 (2021).

45. Jones, S. & Thornton, J. M. Principles of protein-protein interactions. Proc. Natl. Acad. Sci. U.S.A. 93, 13–20 (1996).

46. Chothia, C. & Janin, J. Principles of protein–protein recognition. Nature 256, 705–708 (1975).

47. Ilgü, H. et al. High-resolution structure of the amino acid transporter AdiC reveals insights into the role of water molecules and networks in oligomerization and substrate binding. BMC Biol 19, 179 (2021).

48. Blount, K. F., Wang, J. X., Lim, J., Sudarsan, N. & Breaker, R. R. Antibacterial lysine analogs that target lysine riboswitches. Nat Chem Biol 3, 44–49 (2007).

49. Ataide, S. F. et al. Mechanisms of Resistance to an Amino Acid Antibiotic That Targets Translation. ACS Chem. Biol. 2, 819–827 (2007).

50. Jester, B. C., Levengood, J. D., Roy, H., Ibba, M. & Devine, K. M. Nonorthologous replacement of lysyl-tRNA synthetase prevents addition of lysine analogues to the genetic code. Proc. Natl. Acad. Sci. U.S.A. 100, 14351–14356 (2003).

51. Di Girolamo, M., Busiello, V., Cini, C., Foppoli, C. & De Marco, C. Thialysine utilization by E. coli and its effects on cell growth. Mol Cell Biochem 46, 43–48 (1982).

52. Hirshfield, I. N. & Zamecnik, P. C. Thiosine-resistant mutants of *Escherichia coli* K-12 with growth-medium-dependent lysl-tRNA synthetase activity. I. Isolation and physiological characterization. Biochim Biophys Acta 259, 330–343 (1972).

53. Hirshfield, I. N., Tomford, J. W. & Zamecnik, P. C. Thiosine-resistant mutants of *Escherichia coli* K-12 with growth-medium-dependent lysyl-tRNA synthetase activity.II. Evidence for an altered lysyl-tRNA synthetase. Biochim Biophys Acta 259, 344–356 (1972).

54. Di Girolamo, M., Coccia, R., Blarzino, C., Di Girolamo, A. & Busiello, V. Degradation of thialysine- or selenalysine-containing abnormal proteins in E. coli. Biochem Int 16, 1033–1040 (1988).

55. Drew, D., Lerch, M., Kunji, E., Slotboom, D.-J. & de Gier, J.-W. Optimization of membrane protein overexpression and purification using GFP fusions. Nat Methods 3, 303–313 (2006).

56. Lobley, A., Whitmore, L. & Wallace, B. A. DICHROWEB: an interactive website for the analysis of protein secondary structure from circular dichroism spectra. Bioinformatics 18, 211–212 (2002).

57. Whitmore, L. & Wallace, B. A. DICHROWEB, an online server for protein secondary structure analyses from circular dichroism spectroscopic data. Nucleic Acids Res. 32, W668–673 (2004).

58. Whitmore, L. & Wallace, B. A. Protein secondary structure analyses from circular dichroism spectroscopy: methods and reference databases. Biopolymers 89, 392–400 (2008).

59. Jerabek-Willemsen, M., Wienken, C. J., Braun, D., Baaske, P. & Duhr, S. Molecular interaction studies using microscale thermophoresis. Assay Drug Dev. Technol. 9, 342–353 (2011).

60. Wienken, C. J., Baaske, P., Rothbauer, U., Braun, D. & Duhr, S. Protein-binding assays in biological liquids using microscale thermophoresis. Nat. Commun. 1, 100 (2010).

61. Ward, A. et al. The Amplified Expression Identification, Purification, and Properties of Hexahistidine-Tagged Bacterial Membrane Transport Proteins. In ‘Membrane Transport - a Practical Approach’. Baldwin, S.A. (Ed.) Chapter 6, Pp141-166. (Blackwell’s Press: Oxford UK, 2000).

62. Schorb, M., Haberbosch, I., Hagen, W. J. H., Schwab, Y. & Mastronarde, D. N. Software tools for automated transmission electron microscopy. Nat Methods 16, 471–477 (2019).

63. Punjani, A., Rubinstein, J. L., Fleet, D. J. & Brubaker, M. A. cryoSPARC: algorithms for rapid unsupervised cryo-EM structure determination. Nat Methods 14, 290–296 (2017).

64. Jumper, J. et al. Highly accurate protein structure prediction with AlphaFold. Nature 596, 583–589 (2021).

65. Emsley, P., Lohkamp, B., Scott, W. G. & Cowtan, K. Features and development of Coot. Acta Crystallogr D Biol Crystallogr 66, 486–501 (2010).

66. Liebschner, D. et al. Macromolecular structure determination using X-rays, neutrons and electrons: recent developments in *Phenix*. Acta Crystallogr D Struct Biol 75, 861–877 (2019).

67. The PyMOL Molecular Graphics System, Version 3.0 Schrödinger, LLC.

68. Pettersen, E. F. et al. UCSF Chimera—A visualization system for exploratory research and analysis. J Comput Chem 25, 1605–1612 (2004).

69. Goddard, T. D. et al. UCSF ChimeraX: Meeting modern challenges in visualization and analysis. Protein Science 27, 14–25 (2018).

70. Pardon, E. et al. A general protocol for the generation of Nanobodies for structural biology. Nat Protoc 9, 674–693 (2014).

71. Nji, E., Chatzikyriakidou, Y., Landreh, M. & Drew, D. An engineered thermal-shift screen reveals specific lipid preferences of eukaryotic and prokaryotic membrane proteins. Nat Commun 9, 4253 (2018).

72. Chatzikyriakidou, Y., Ahn, D.-H., Nji, E. & Drew, D. The GFP thermal shift assay for screening ligand and lipid interactions to solute carrier transporters. Nat Protoc 16, 5357–5376 (2021).

73. Madhavi Sastry, G., Adzhigirey, M., Day, T., Annabhimoju, R. & Sherman, W. Protein and ligand preparation: parameters, protocols, and influence on virtual screening enrichments. J Comput Aided Mol Des 27, 221–234 (2013).

74. Lu, C. et al. OPLS4: Improving Force Field Accuracy on Challenging Regimes of Chemical Space. J Chem Theory Comput 17, 4291–4300 (2021).

75. Friesner, R. A. et al. Glide: a new approach for rapid, accurate docking and scoring. 1. Method and assessment of docking accuracy. J Med Chem 47, 1739–1749 (2004).

76. Sievers, F. & Higgins, D. G. Clustal Omega for making accurate alignments of many protein sequences. Protein Sci 27, 135–145 (2018).

