## Supplementary materials for "Structural insights into *Pseudomonas aeruginosa* lysine-specific uptake mechanism for extremely low pH regulation"

Fig. S1.

**a**

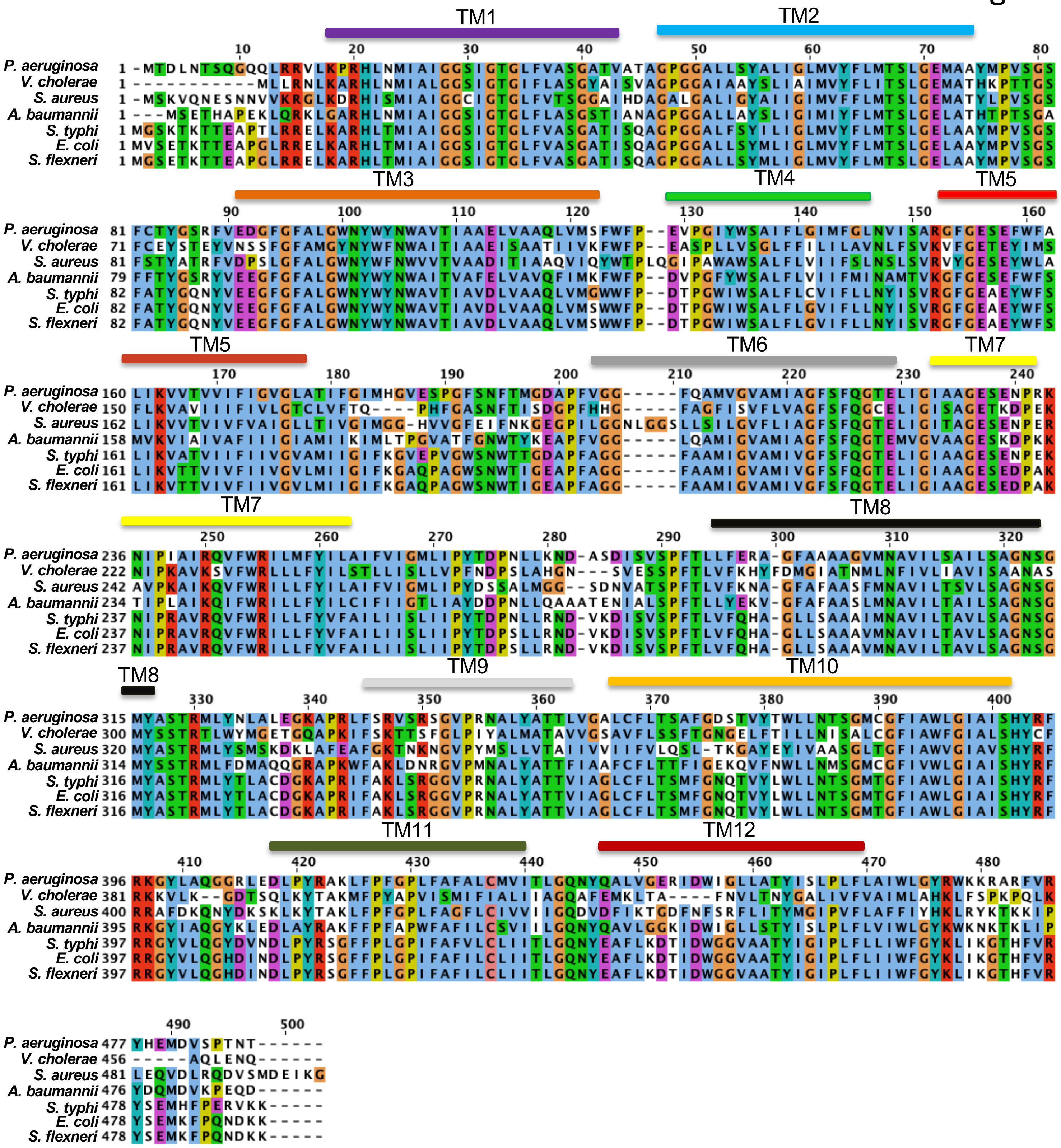


**b**

| LysP | *P. aeruginosa* | *V. cholerae* | *S. aureus* | *A. baumannii* | *S. typhi* | *E. coli* | *S. flexneri* |
|---|---|---|---|---|---|---|---|
| ***P. aeruginosa*** | 100 | | | | | | |
| ***V. cholerae*** | 45.9 | 100 | | | | | |
| ***S. aureus*** | 51.5 | 42.8 | 100 | | | | |
| ***A. baumannii*** | 65.8 | 46.3 | 48.3 | 100 | | | |
| ***S. typhi*** | 70.8 | 46.7 | 50.1 | 60.8 | 100 | | |
| ***E. coli*** | 70.2 | 46.7 | 49.7 | 61.2 | 94.5 | 100 | |
| ***S. flexneri*** | 70.2 | 46.7 | 49.7 | 61.2 | 94.7 | 99.8 | 100 |

Fig. S2.

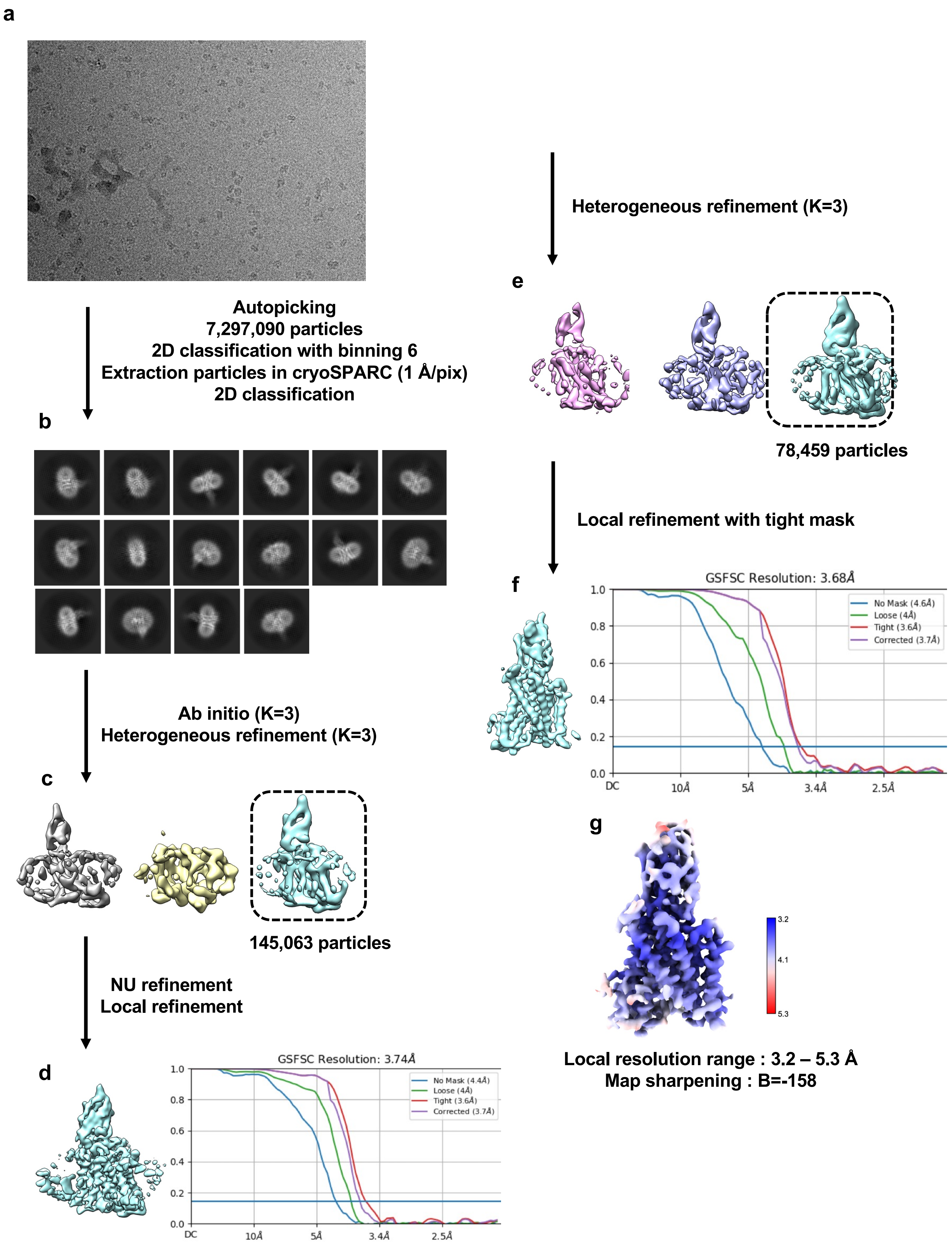
a
Autopicking
7,297,090 particles
2D classification with binning 6
Extraction particles in cryoSPARC (1 Å/pix)
2D classification
b
Ab initio (K=3)
Heterogeneous refinement (K=3)
c
145,063 particles
NU refinement
Local refinement
d
GSFSC Resolution: 3.74Å
No Mask (4.4Å)
Loose (4Å)
Tight (3.6Å)
Corrected (3.7Å)
1.0
0.8
0.6
0.4
0.2
0.0
DC
10Å
5Å
3.4Å
2.5Å
Heterogeneous refinement (K=3)
e
78,459 particles
Local refinement with tight mask
f
GSFSC Resolution: 3.68Å
No Mask (4.6Å)
Loose (4Å)
Tight (3.6Å)
Corrected (3.7Å)
1.0
0.8
0.6
0.4
0.2
0.0
DC
10Å
5Å
3.4Å
2.5Å
g
3.2
4.1
5.3
Local resolution range : 3.2 – 5.3 Å
Map sharpening : B=-158

Fig. S3.

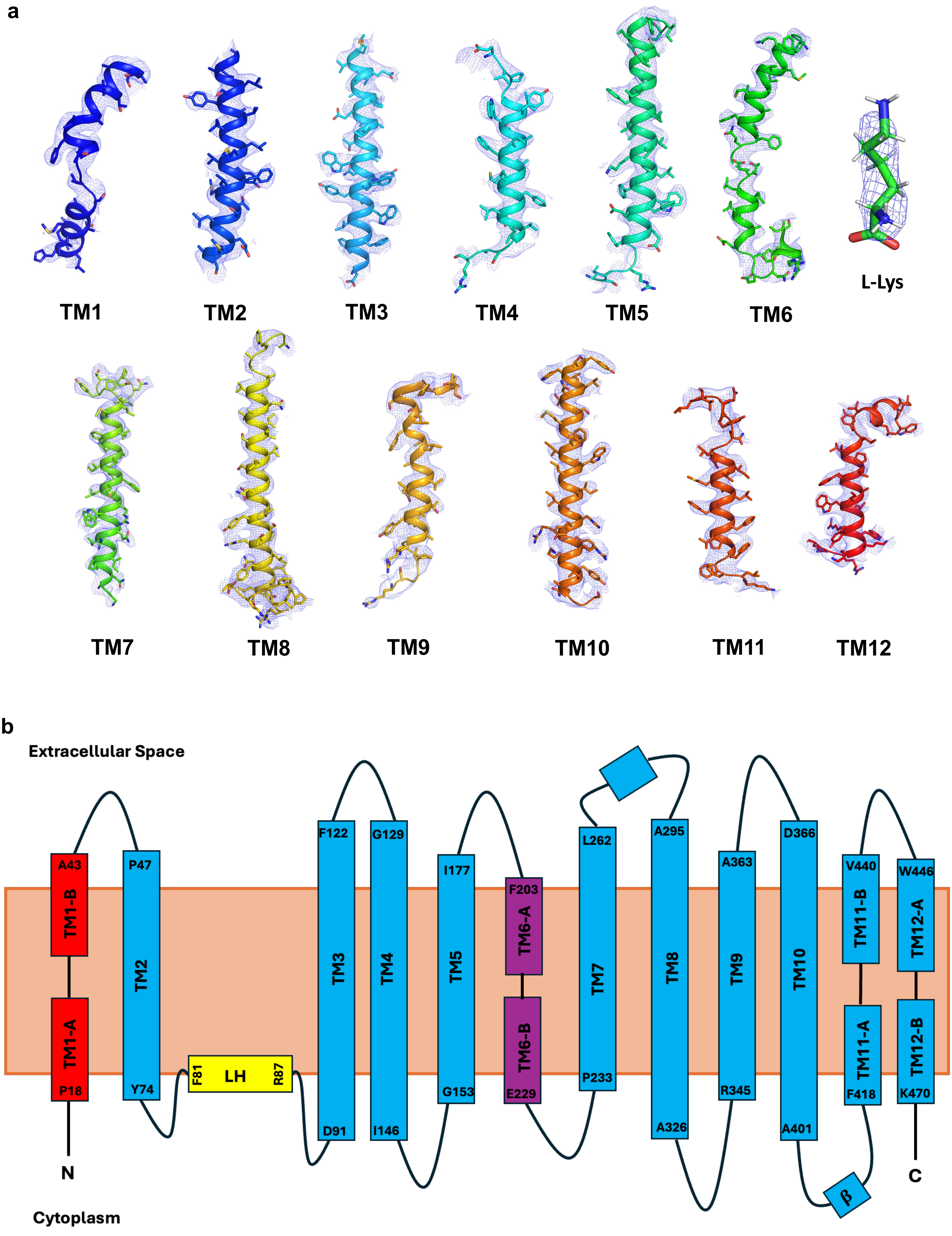
a
TM1
TM2
TM3
TM4
TM5
TM6
L-Lys
TM7
TM8
TM9
TM10
TM11
TM12
b
Extracellular Space
Cytoplasm
N
C
A43
TM1-B
TM1-A
P18
P47
TM2
Y74
F81
LH
R87
F122
TM3
D91
G129
TM4
I146
I177
TM5
G153
F203
TM6-A
TM6-B
E229
L262
TM7
P233
A295
TM8
A326
A363
TM9
R345
D366
TM10
A401
V440
TM11-B
TM11-A
F418
W446
TM12-A
TM12-B
K470
β

Fig. S4.

a

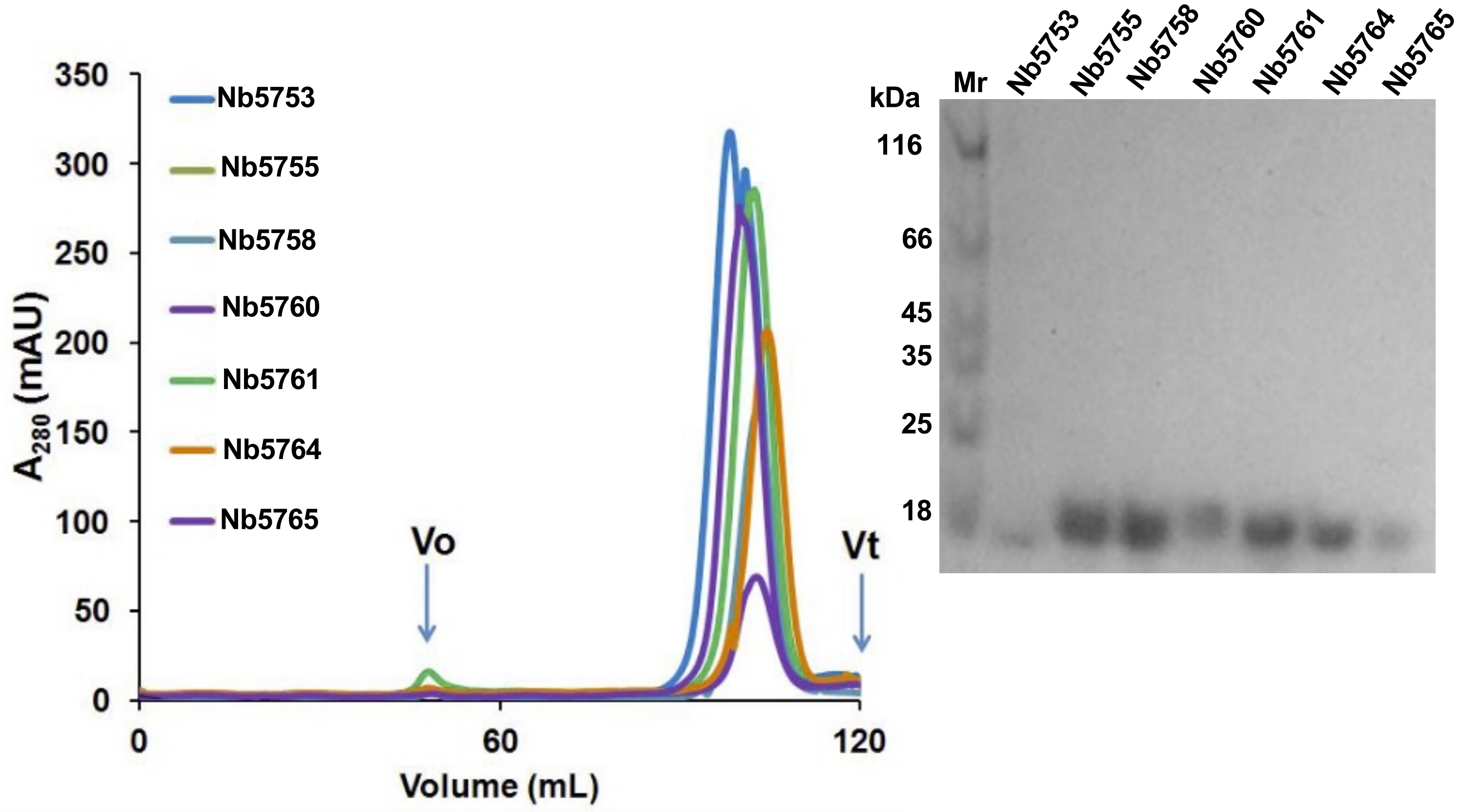
Nb5753
Nb5755
Nb5758
Nb5760
Nb5761
Nb5764
Nb5765
Vo
Vt
A280 (mAU)
Volume (mL)
0
60
120
350
300
250
200
150
100
50
kDa
Mr
116
66
45
35
25
18

b

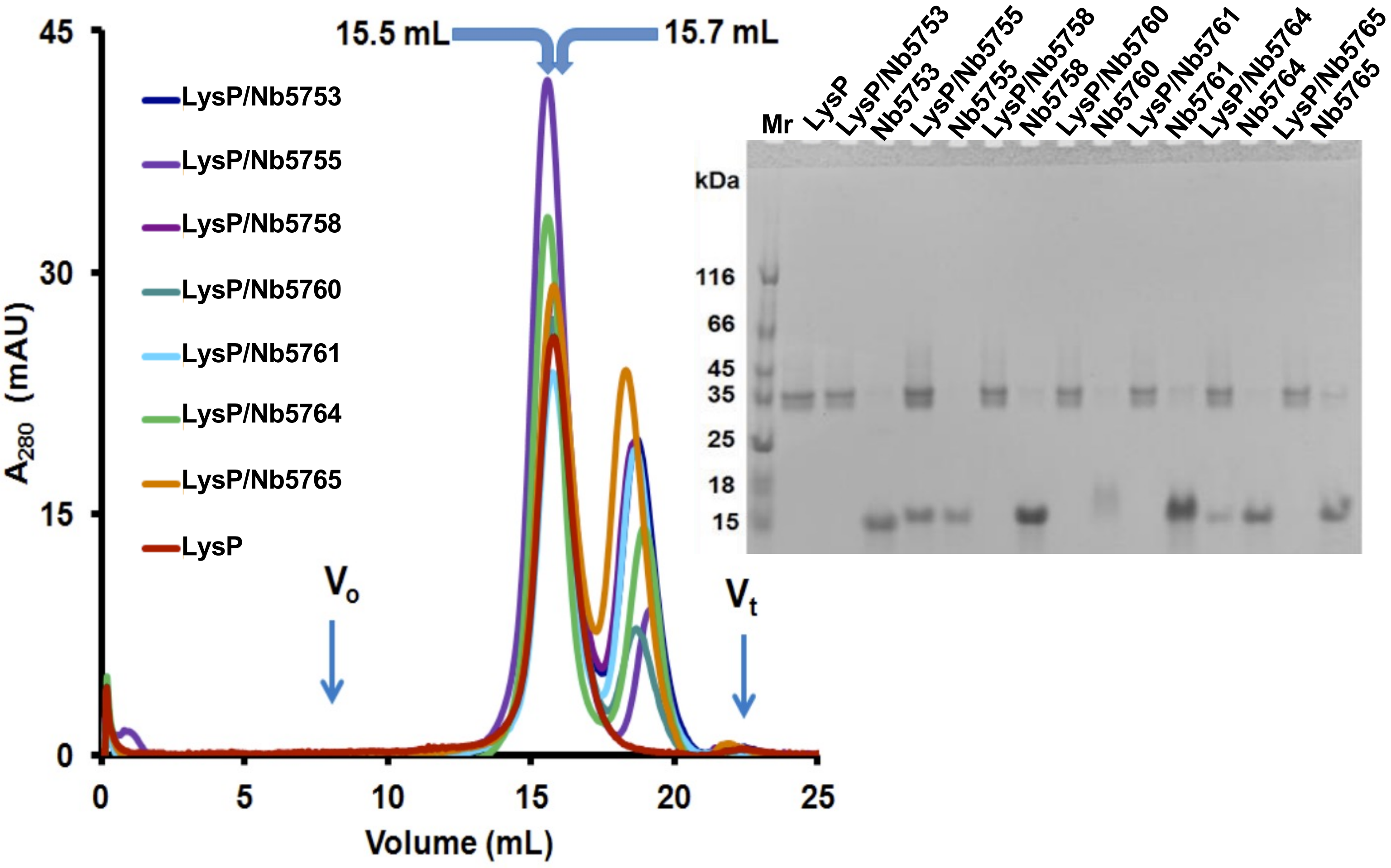
15.5 mL
15.7 mL
LysP/Nb5753
LysP/Nb5755
LysP/Nb5758
LysP/Nb5760
LysP/Nb5761
LysP/Nb5764
LysP/Nb5765
LysP
Vo
Vt
A280 (mAU)
Volume (mL)
0
5
10
15
20
25
30
45
Mr
LysP
kDa
116
66
45
35
25
18
15

Fig. S5.

a

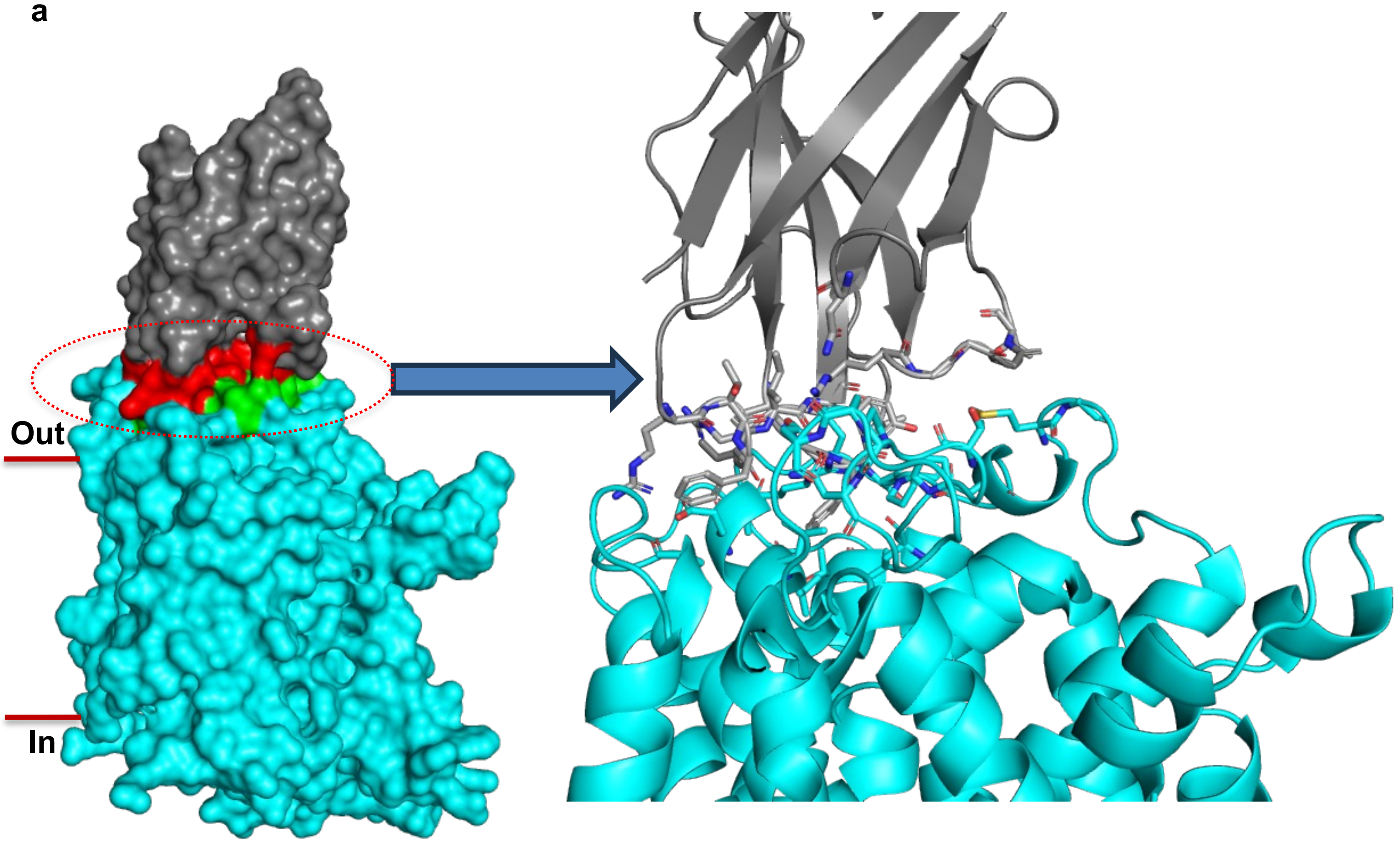
Out
In

b

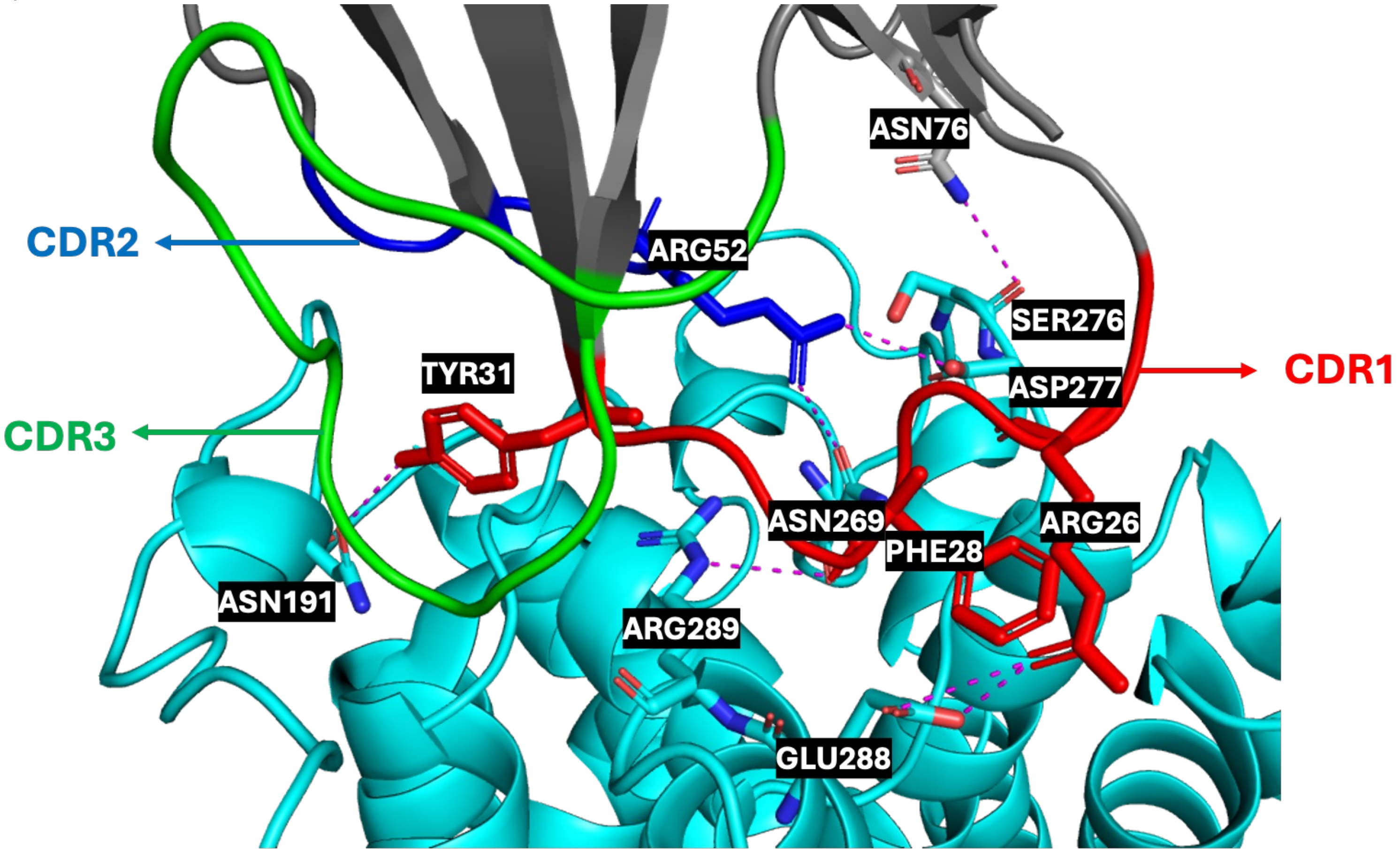
ASN76
CDR2
ARG52
SER276
TYR31
ASP277
CDR1
CDR3
ASN269
ARG26
PHE28
ASN191
ARG289
GLU288

Fig. S6.

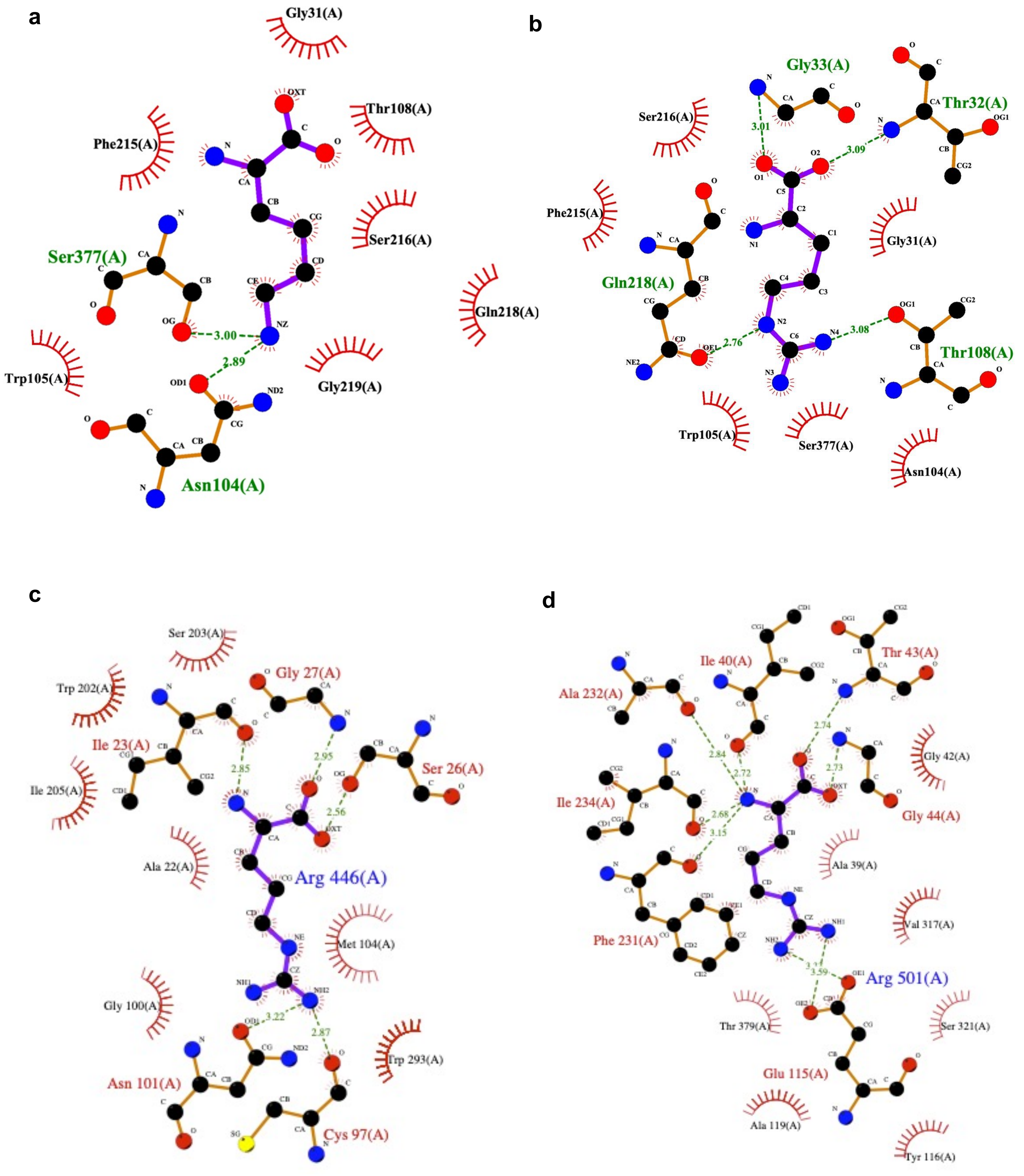
a
Gly31(A)
Thr108(A)
Phe215(A)
Ser216(A)
Ser377(A)
Gln218(A)
Trp105(A)
Gly219(A)
Asn104(A)
3.00
2.89
b
Gly33(A)
Thr32(A)
Ser216(A)
Phe215(A)
Gly31(A)
Gln218(A)
Thr108(A)
Trp105(A)
Ser377(A)
Asn104(A)
3.01
3.09
2.76
3.08
c
Ser 203(A)
Trp 202(A)
Gly 27(A)
Ile 23(A)
Ser 26(A)
Ile 205(A)
Ala 22(A)
Arg 446(A)
Met 104(A)
Gly 100(A)
Trp 293(A)
Asn 101(A)
Cys 97(A)
2.85
2.95
2.56
3.22
2.87
d
Ile 40(A)
Thr 43(A)
Ala 232(A)
Gly 42(A)
Ile 234(A)
Gly 44(A)
Ala 39(A)
Val 317(A)
Phe 231(A)
Arg 501(A)
Thr 379(A)
Ser 321(A)
Glu 115(A)
Ala 119(A)
Tyr 116(A)
2.84
2.72
2.74
2.73
2.68
3.15
3.59

Fig. S7.

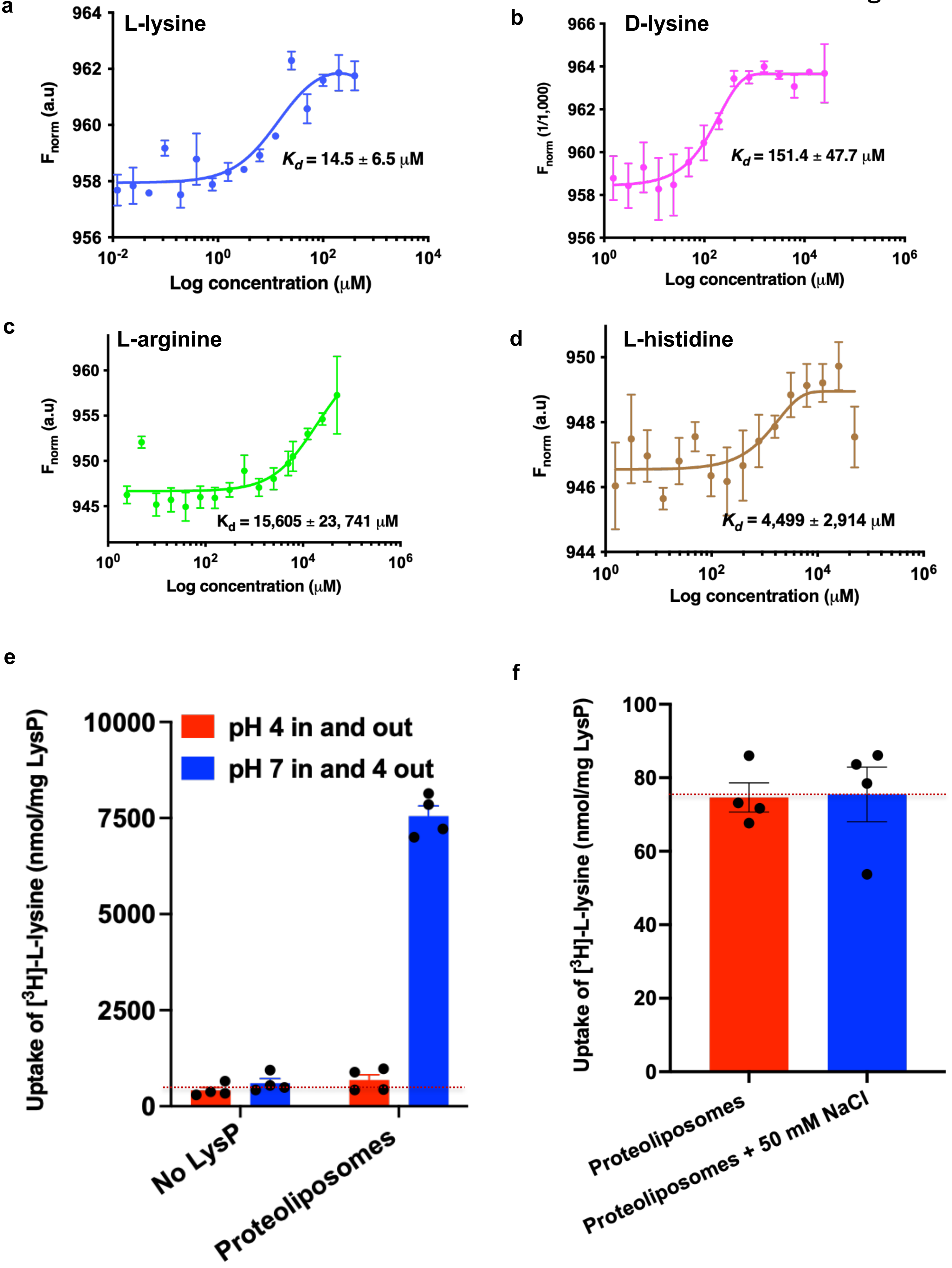
a
L-lysine
$F_{norm}$ (a.u)
$K_d$ = 14.5 ± 6.5 μM
Log concentration (μM)
b
D-lysine
$F_{norm}$ (1/1,000)
$K_d$ = 151.4 ± 47.7 μM
Log concentration (μM)
c
L-arginine
$F_{norm}$ (a.u)
$K_d$ = 15,605 ± 23, 741 μM
Log concentration (μM)
d
L-histidine
$F_{norm}$ (a.u)
$K_d$ = 4,499 ± 2,914 μM
Log concentration (μM)
e
Uptake of [$^3$H]-L-lysine (nmol/mg LysP)
pH 4 in and out
pH 7 in and 4 out
No LysP
Proteoliposomes
f
Uptake of [$^3$H]-L-lysine (nmol/mg LysP)
Proteoliposomes
Proteoliposomes + 50 mM NaCl

Fig. S8.

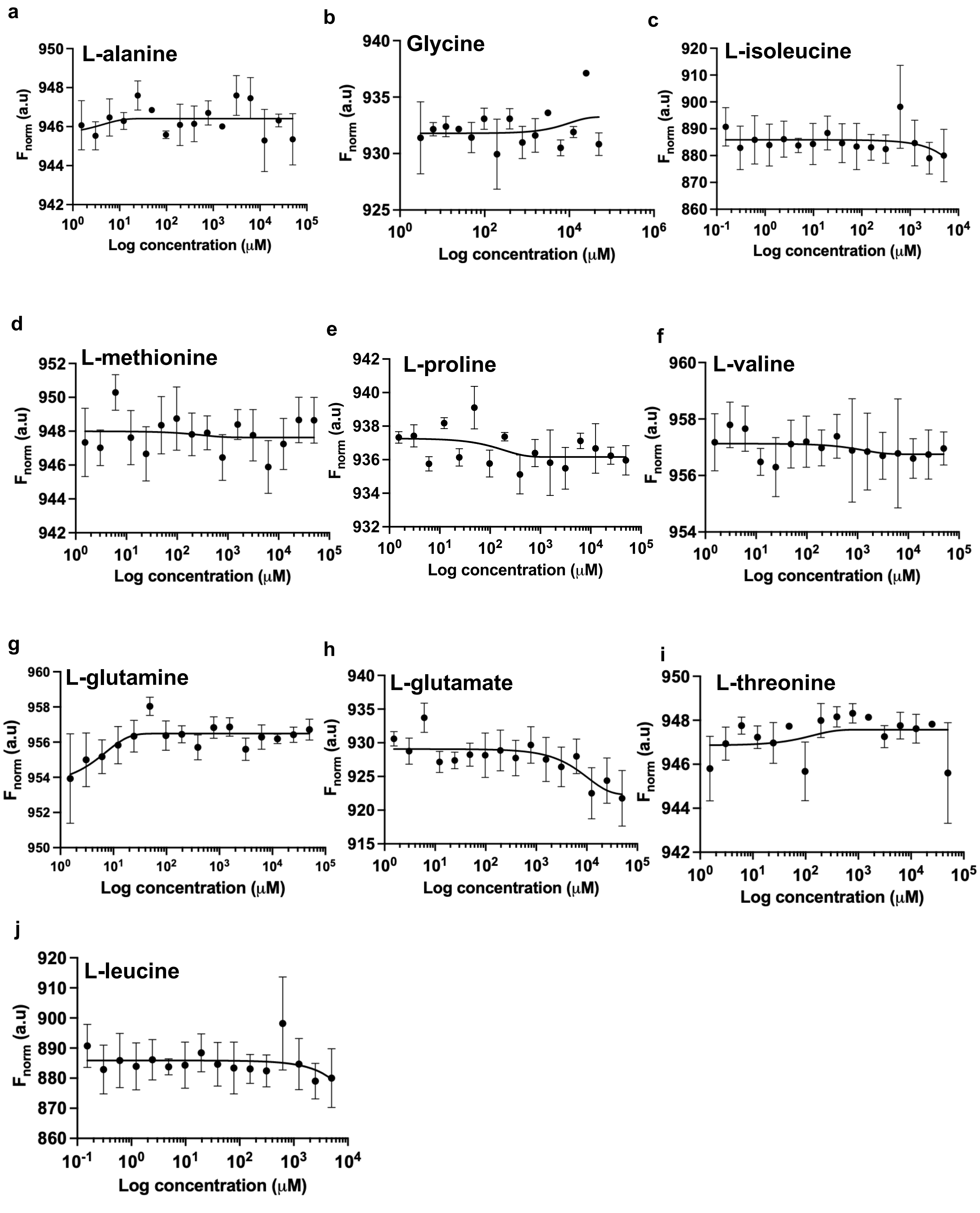

a
L-alanine
$F_{norm}$ (a.u)
Log concentration (μM)
b
Glycine
$F_{norm}$ (a.u)
Log concentration (μM)
c
L-isoleucine
$F_{norm}$ (a.u)
Log concentration (μM)
d
L-methionine
$F_{norm}$ (a.u)
Log concentration (μM)
e
L-proline
$F_{norm}$ (a.u)
Log concentration (μM)
f
L-valine
$F_{norm}$ (a.u)
Log concentration (μM)
g
L-glutamine
$F_{norm}$ (a.u)
Log concentration (μM)
h
L-glutamate
$F_{norm}$ (a.u)
Log concentration (μM)
i
L-threonine
$F_{norm}$ (a.u)
Log concentration (μM)
j
L-leucine
$F_{norm}$ (a.u)
Log concentration (μM)
